# POLQ-driven repair scars shape the immunogenic landscape of homologous recombination-deficient pancreatic cancer

**DOI:** 10.64898/2026.03.15.711961

**Authors:** Wungki Park, Shigeaki Umeda, Marc Hilmi, Catherine O’Connor, Roshan Sharma, Nuray Tezcan, Haochen Zhang, Yingjie Zhu, Carly Schwartz, Amin Yaqubie, Anna Varghese, Kevin Soares, Vaia Florou, Daewon Kim, Steven Maron, Guillem A. Martinez, Fiyinfolu Balogun, Caitlin McIntyre, Daehee Kim, Kenneth H. Yu, Joanne F. Chou, Akimasa Hayashi, Fergus Keane, Danny Khalil, Walid K Chatila, Marinela Capanu, Ronan Chaligne, Michael J Pishvaian, Chaitanya Bandlamudi, Nicolas Lecomte, Michael Berger, Olca Basturk, Vinod Balachandran, Dana Pe’er, Benoit Rousseau, Benjamin Greenbaum, Agnel Sfeir, Christine Iacobuzio-Donahue, Nadeem Riaz, Eileen M. O’Reilly

## Abstract

Pancreatic cancer (PC) is broadly resistant to immune checkpoint blockade, although a subset of homologous recombination-deficient (HRD) tumors exhibits durable immune engagement. The genomic features that distinguish these immune-responsive tumors from immune-inert HRD tumors remain poorly understood. Here we identify a microhomology-mediated end joining (MMEJ) repair scar, the MMEJ Deletion Footprint (MDF), as a genomic readout of POLQ-associated error-prone repair that enriches for frameshift indels. Across the multi-omic discovery cohort integrating tumor genomics, single-nucleus transcriptomics and spatial immune profiling, MDF-high HRD PC exhibited increased frameshift-indel-derived neoantigens and interferon programs. MDF was further associated with remodeling of the myeloid compartment toward MHC II-high dendritic cell-like antigen-presenting macrophage states and the immune synapse architecture marked by increased spatial interaction between APC-like macrophages and cytotoxic CD8^+^ T cells. These tissue-level features aligned with a functional trajectory shift of CD8^+^ T cells, consistent with effective anti-tumor immunity and was associated with favorable clinical outcomes of patients. Together, our findings position MMEJ-linked repair scarring as actionable biology that connects an HRD genotype to immune organization and suggests rational immunotherapy combinations that may enhance antigen presentation and myeloid activation to extend durable benefit in HRD-lineage cancers.

## INTRODUCTION

Immunotherapy has transformed treatment for many solid tumors, but its impact in pancreatic cancer (PC) remains limited^1,2^. Most patients with advanced PC do not benefit from immune checkpoint blockade (ICB). Clinical trials evaluating programmed cell death 1 (PD-1) and cytotoxic T-lymphocyte associated protein-4 (CTLA4) inhibition in unselected PC have yielded disappointing results^3–6^. Yet, a small subset of patients, often those with underlying DNA damage repair deficiencies (DDR), experience durable clinical benefit, suggesting a distinct biological subset amenable to immunotherapy^7,8^.

The most definitively immunogenic subset of PC comprises the rare (∼1%) microsatellite instability-high (MSI-H) tumors with mismatch repair deficiency (dMMR)^7,9^. These tumors harbor high-quality neoantigens, largely driven by frameshift indels and missense mutations, which promote robust neoantigen-cognate immune cell infiltration at baseline and enable durable responses to ICB^9–13^. Similarly, homologous recombination deficient (HRD; 5-8%) tumors harbor *BRCA1/2*, *PALB2, or RAD51C/D* mutations have been observed to have increased indel burden and immune infiltration^14–18^. However, not all HRD tumors are equally immunogenic and predictive biomarkers and tumor microenvironmental features that stratify immune responsiveness within HRD PC remain undefined.

In the POLAR trial, we recently evaluated a biomarker-driven precision immunotherapy maintenance strategy in patients with platinum-sensitive metastatic PC, with or without HRD^19^. This phase 2 trial investigated the combination of pembrolizumab (anti-PD-1) plus olaparib (poly-ADP-ribosyl polymerase inhibitor; PARPi) and demonstrated an exploratory response rate of 53% and a 6-month progression-free survival (PFS) rate of 64% (95% CI: 49-82). Remarkably, the 3-year overall survival (OS) rate was 44% (95% CI: 28-69) and this is an exceptional outcome in this notoriously immunotherapy-resistant metastatic cancer.

Homologous recombination deficiency creates a state of impaired high-fidelity double-strand break (DSB) repair, forcing malignant cells to rely on alternative, error-prone repair pathways to maintain genome integrity^20^. One such pathway is microhomology-mediated end joining (MMEJ), a POLQ-dependent DSB repair mechanism that utilizes short microhomology sequences to rejoin broken DNA ends. Unlike homologous recombination, MMEJ is intrinsically mutagenic and generates characteristic deletions near microhomology sites, leaving distinct genomic scars^21–23^. Although MMEJ has been mechanistically characterized and implicated in HRD tumor evolution, its contribution to shaping immunogenic mutational landscapes and tumor-immune interactions in human PC remains undefined. We therefore hypothesized that POLQ-mediated MMEJ repair scarring constitutes a genomic substrate for immunogenicity in HRD PC.

Here, we define a MMEJ Deletion Footprint (MDF), a quantitative metric derived from whole-exome sequencing that captures the abundance of short deletions within the characteristic 6–20 base pair range associated with POLQ-mediated MMEJ repair in human PC. Using multi-omic discovery and validation datasets, we show that MDF-high HRD PC harbor increased high-quality neoantigen burden and a favorable tumor immune microenvironment characterized by enriched cytotoxic T cells and reduced immunosuppressive Tumor-Associated Macrophages (TAMs). These tissue-level features provide a mechanistic framework linking HRD genotype to immune organization and help explain the improved clinical outcomes observed in select patients treated in the POLAR trial. Together, these findings establish MDF as a genomic biomarker of POLQ-driven immunogenicity and support precision immunotherapy strategies for HRD-lineage cancers beyond PC.

## RESULTS

### Distinct DNA repair–deficient subgroups define immunogenic pancreatic cancer

To establish the molecular basis of immunogenic PC, we assembled a multiomic discovery cohort of 71 patients with metastatic PC (immunogenic PC; iPC)^24^. Whole-exome sequencing was performed in all patients (WES, N=71), and bulk RNA-seq was available for 56 patients and single-nucleus RNA sequencing (snRNA-seq) for 23 patients (**Figure S1**). Tumors were stratified into mismatch repair deficient (dMMR, N=7, 10%), HRD (*BRCA1 or BRCA2* mutations, N=18, 25%) and non-core HRD (*ATM, CHEK2, FANCA or RAD50* mutations, N=7, 10%) subgroups, reflecting biologically distinct mechanisms of genomic instability across DNA damage repair (DDR)-defined classes (**Figure 1A; Figure S1-S3; Table S1**). The remaining tumors were classified as homologous recombination-proficient (HRP, N=39, 55%).

**Figure 1.**
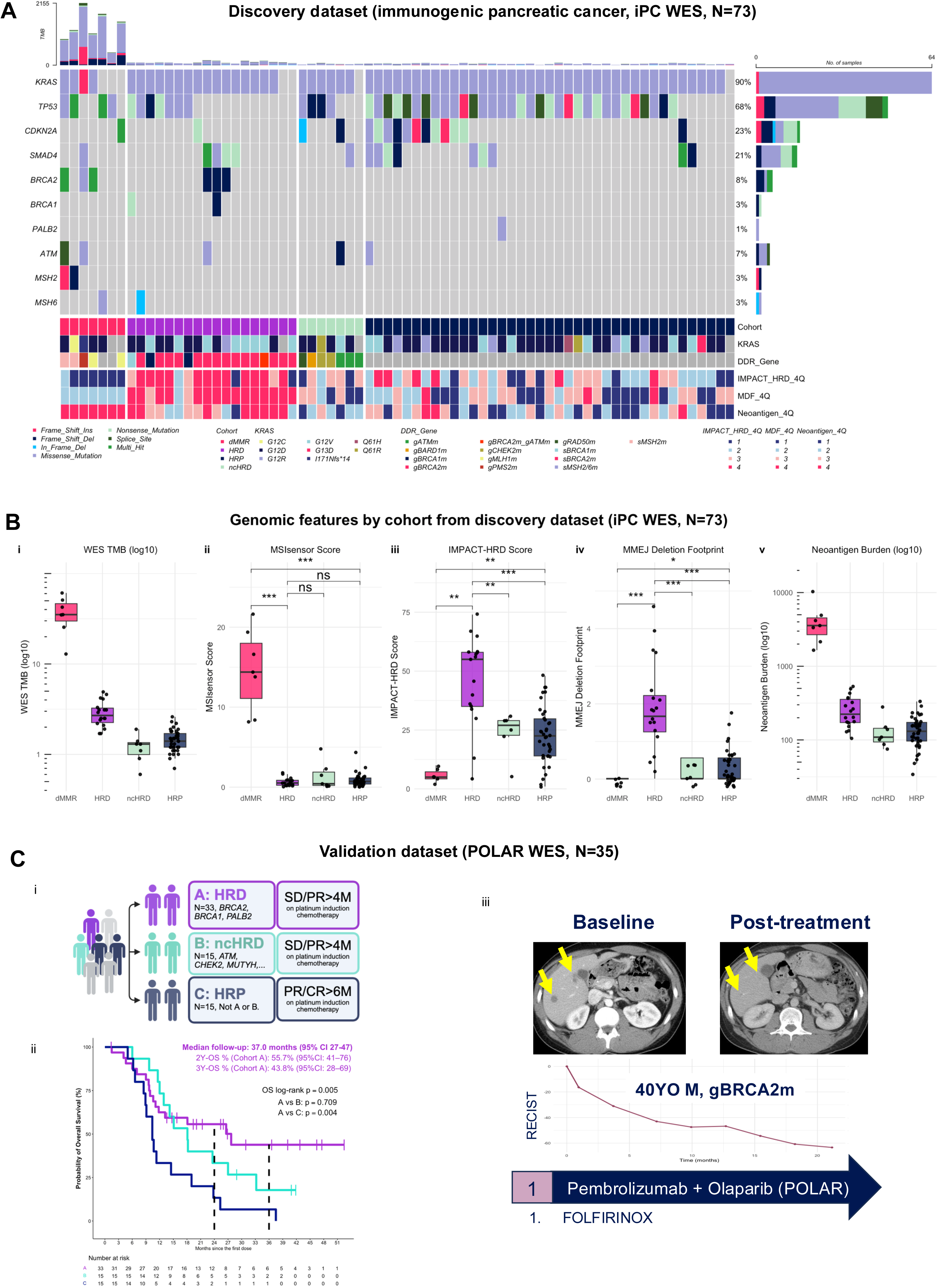
Distinct DNA damage repair deficient subgroups define immunogenic pancreatic cancer. **(A)** Oncoprint of the discovery dataset (immunogenic pancreatic cancer cohort [iPC], WES, N=73) showing somatic alterations in DNA damage repair (DDR) genes (patients included all had germline *BRCA1*, *BRCA2, or PALB2* mutation). Samples are stratified into mismatch repair deficient (dMMR, red), homologous recombination deficient (HRD, purple), non-core HRD (ncHRD, green), and homologous recombination proficient (HRP, navy) subgroups. Annotations include *KRAS* status, DDR gene alterations, IMPACT-HRD score quartile (4Q), MMEJ deletion footprint (MDF) quartile (4Q), and neoantigen burden quartile (4Q). **(B)** Genomic features across subgroups. Panels show: (i) WES-derived tumor mutation burden (log10), (ii) MSIsensor score, and (iii) IMPACT-HRD score, (iv) MDF (MMEJ Deletion Footprint) and (v) Neoantigen Burden (log10). **(C)** Validation in the POLAR trial dataset (POLAR WES, N=35). (i) POLAR cohort definition schema based on DNA repair deficient status and clinical response to platinum-based chemotherapy. (ii) Kaplan-Meier overall survival (OS) curves showing durable outcomes in HRD pancreatic cancer (Cohort A, purple) compared with non-core HRD (Cohort B) and homologous recombination proficient (HRP, HRD-negative, Cohort C) tumors. (iii) Representative radiographic response in a patient with germline *BRCA2*-mutant pancreatic cancer treated with pembrolizumab plus olaparib maintenance following first-line platinum chemotherapy. **Supplementary Reference:** See also Figures S1–S3 and Table S1. **Abbreviations:** PC, pancreatic cancer; dMMR, mismatch repair deficient; HRD, homologous recombination deficient; ncHRD, non-core HRD; HRP, homologous recombination proficient; TMB, tumor mutational burden; ICB, immune checkpoint blockade; MMEJ, microhomology-mediated end joining; MDF, MMEJ deletion footprint.

Comprehensive WES profiling in the discovery cohort revealed marked differences in genomic features across DDR subgroups (**Figure 1B**). As expected, dMMR tumors exhibited the highest tumor mutational burden (TMB), MSIsensor scores and neoantigen burden. By contrast, HRD tumors demonstrated only modest increases in TMB, but were significantly enriched for higher IMPACT-HRD scores and microhomology-mediated end joining (MMEJ) Deletion Footprint (MDF) scores, distinguishing them from both ncHRD and HRP tumors. These findings suggest an alternative, non-TMB-driven pathway for the immunogenicity in HRD PC.

Clinically, dMMR tumors define a canonically immunogenic subset with sensitivity to immune checkpoint blockade (ICB), as illustrated by significantly improved survival among patients receiving ICB compared with those who did not (p=0.0082; **Figure S2A**). Although dMMR tumors are rare in PC, this subgroup demonstrated exceptional responders, exemplified by a patient with a complete response to anti-PD-1 ICB monotherapy and remaining on active surveillance for 2.5 years, accompanied by expansion of circulating CD8⁺ T, CD4⁺ T and CD3⁺CD56⁺CD16⁺ natural killer T cells following treatment (**Figure S2C**). Notably, a subset of HRD tumors from the POLAR trial (Cohort A, **Figure 1C**) also demonstrated durable survival benefit from immunotherapy-based strategies compared to cohort B (ncHRD) and C (HRP) (p=0.005). In the POLAR trial, patients received pembrolizumab and olaparib maintenance as a maintenance after first-line platinum chemotherapy as an induction for more than 4 months. Additionally, we observed patients with germline BRCA1 or BRCA2 mutations had exceptional response to dual immune checkpoint blockade like others (**Figure 2SD)**^8^. These observations raised the central question addressed in this study: why do some HRD tumors exhibit immunogenic features despite a relatively low TMB?

**Figure 2.**
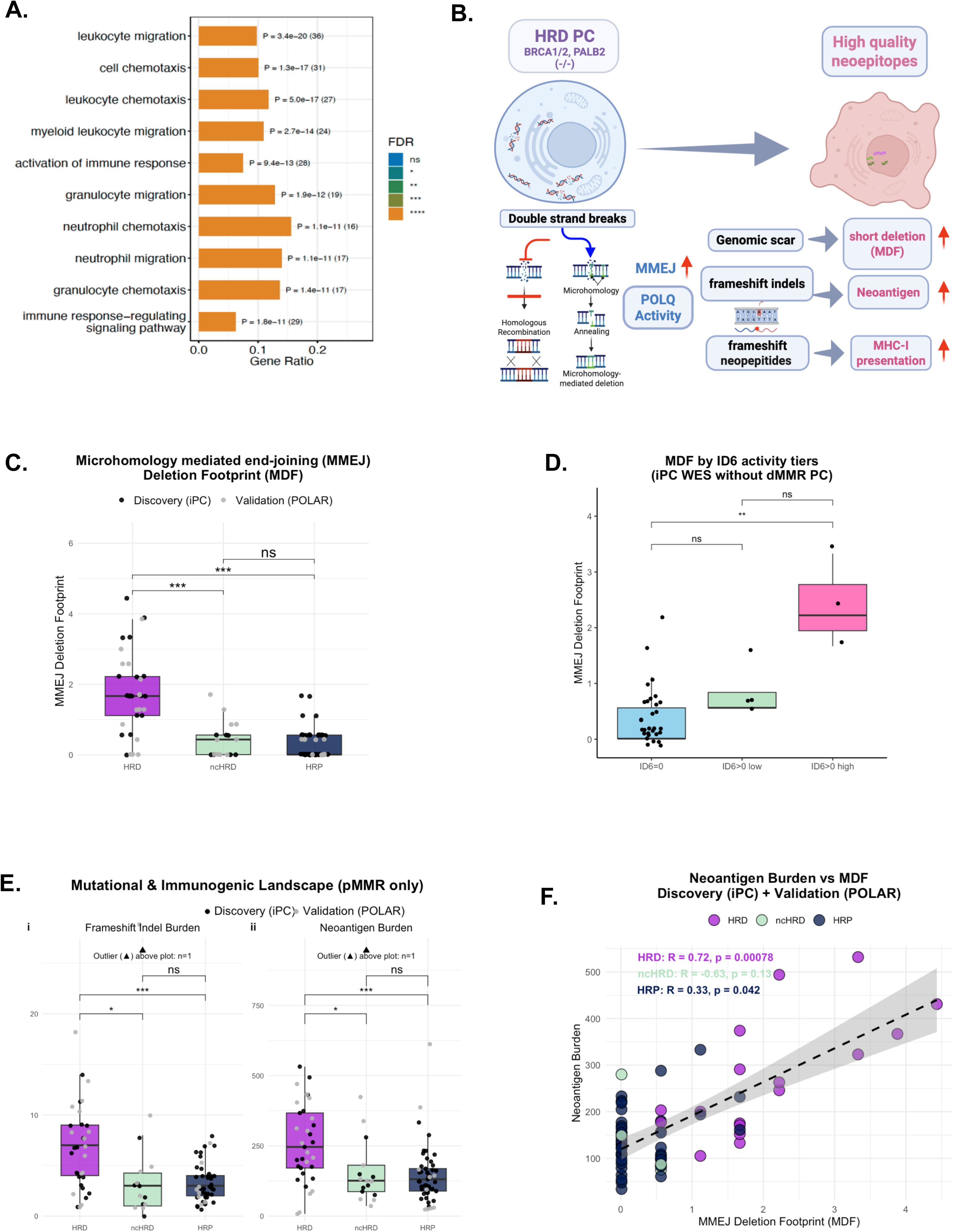
Immunogenicity of HRD PC is associated with DNA repair scar by Microhomology-Mediated End Joining. **(A)** Gene Ontology term enrichment of programs upregulated in HRD PC (N=17) tumors compared to HRP (N=39) tumors highlights immune activation, leukocyte migration, and chemotaxis. FDR-adjusted p-values shown. **(B)** Schema for immunogenicity of HRD PC associated with frameshift indel-derived neoantigens generated by DNA repair of double strand breaks (DSB) via microhomology-mediated end joining **(C)** MMEJ Deletion Footprint (MDF) scores are significantly higher in HRD tumors compared to HRP and ncHRD tumors, supporting DDR subgroup specificity. (Discovery iPC data are black dots and Validation POLAR data are grey dots; ***, p < 0.001 for HRD vs all the other subgroups) **(D)** COSMIC indel mutational signature analysis from the Discovery dataset. ID6 is known to be associated with MMEJ activity. MDF scores were statistically higher in ID6-high samples compared to ID6-undetected samples. **(E)** Mutational and immunogenic landscape in mismatch repair proficient (pMMR, without dMMR) samples. Subplot I shows frameshift indel burden is significantly higher in HRD PC compared to ncHRD and HRP PC samples. Same for neoantigen burden in same pattern. (Discovery iPC data are black dots and Validation POLAR data are grey dots; ***, p < 0.001 for HRD vs all the other subgroups) **(F)** The association of neoantigen burden and MDF was plotted per each cohort from combined cohorts. Pearson’s analysis show that in HRD PC has the strongest statistically significant correlation between neoantigen burden and MDF (R=0.72, p=0.00078) **Supplementary Reference**: See also Figure S3-5 for transcriptional and genomic pattern analyses. **Abbreviations:** DSB, double strand breaks; MMEJ, microhomology-mediated end joining; MDF, MMEJ Deletion Footprint; iPC, immunogenic pancreatic cancer; HRD, homologous recombination–deficient; ncHRD, non-core HRD; HRP, homologous recombination–proficient; dMMR, mismatch repair–deficient; ID6, COSMIC indel signature 6

To further contextualize these findings, we examined canonical oncogenic drivers in the discovery cohort across all groups. Mutations in *KRAS*, *TP53* and *CDKN2A* were broadly similar across cohorts. *KRAS* mutations were present in 5/7 dMMR PC, 17/18 HRD and 38/39 HRP tumors, with similar distributions of hotspot alleles (G12D, G12V, G12R and Q61X) across groups (**Figure 1A; Table S1B**). In contrast, *KRAS* wild-type tumors and non-hotspot mutations including *KRAS* I171Nfs*14, were observed primarily in dMMR PC, consistent with the indel-rich mutational processes of mismatch repair deficiency. Importantly, HRD tumors largely retained canonical *KRAS* mutations, indicating that immunogenic HRD PC arise within the typical *KRAS*-driven pancreatic tumorigenic framework.

Within HRD tumors, germline *BRCA2* mutations were most frequent (N=15), followed by germline *BRCA1* mutations (N=3). ncHRD tumors were enriched for germline *ATM, CHEK2*, *RAD50* and *BARD1* mutations. Biallelic inactivation, defined by co-occurrence of pathogenic mutations and locus-specific loss-of-heterozygosity (LOH), was more frequent in canonical driver genes of HR genes within HRD tumors (*BRCA1/2,* 89%) compared with ncHRD tumors (57%) (**Table S1B**).

Taken together, these findings indicate that immunogenic PC may be enriched within two biologically distinct DDR subgroups, dMMR (1%) and HRD (∼8%), each arising within a characteristic genomic landscape (**Figure S3**). Because dMMR tumors represent a well-established immunogenic subtype driven by hypermutation, subsequent analyses focused on HRD PC, where the mechanisms underlying immunogenicity remain less well understood. This framework enabled us to investigate how POLQ-associated MMEJ repair scarring and MDF contribute to neoantigen generation and tumor-immune interactions in HRD pancreatic cancer.

### Transcriptomic profiling reveals immune activation programs in HRD pancreatic cancer

To determine whether HRD PC exhibits transcriptional evidence of immune engagement, we performed differential expression analysis using bulk RNA-seq data from HRD (N=17) and HRP (N=39) tumors in the discovery cohort. HRD tumors demonstrated robust upregulation of immune-related genes, including those involved in antigen presentation, cytokine signaling and leukocyte trafficking (**Figure S4A-C**).

Gene set enrichment analysis (GSEA) of Hallmark pathways revealed significant enrichment of immune activation programs in HRD tumors, including inflammatory response, interferon-γ signaling, and adaptive immune response pathways. Complementary Gene Ontology enrichment analysis identified activation of immune response, leukocyte migration, chemotaxis, T cell activation and cell-surface receptor signal as the most significantly enriched biological processes in HRD tumors compared to HRP (**Figure 2A; Figure S4D**). These transcriptional programs were absent or attenuated in HRP tumors, establishing that HRD PCs adopt a transcriptionally inflamed phenotype despite lacking the extreme hypermutation characteristic of dMMR tumors.

### HRD tumors are enriched for DNA repair scars quantified by the MMEJ Deletion Footprint

We next sought to identify the genomic mechanisms that might explain immune activation in HRD PC. Fine-scale indel footprint analysis of WES data from both discovery iPC cohort and the independent validation POLAR cohort revealed that HRD tumors were selectively enriched for short deletion events in the 6–20 bp range (**Figure S5A**), a hallmark of microhomology-mediated end joining (MMEJ) repair (**Figure 2B**). To quantify this repair-associated deletion pattern, we derived the MMEJ Deletion Footprint (MDF), a standardized metric based on the abundance of deletions in the 6-20 bp range, which are characteristic of MMEJ repair.

MDF scores were significantly higher in HRD tumors compared with both ncHRD (p<0.001) and HRP (p<0.001) tumors (**Figure 2C**). This enrichment was reproducible across the discovery iPC cohort and independently validated in the POLAR cohort WES, supporting that elevated MMEJ activity is a conserved genomic feature of HRD PC. Further evaluation of COSMIC indel mutational signatures confirmed an enrichment of signature ID6 in HRD tumors, previously known for MMEJ repair activity^25^. MDF scores were significantly higher in ID6-high tumors compared with ID6-undetected tumors (**Figure 2D; Figure S5B**), supporting MDF as a quantitative measure of MMEJ-associated genomic repair scars.

### MMEJ Deletion Footprint generate frameshift indels and high-quality neoantigens

Given the established link between frameshift indels and high-quality neoantigen generation^11,12,26,27^, we next examined whether MDF contribute to tumor immunogenicity. Among mismatch repair–proficient (pMMR) tumors, HRD cancers exhibited significantly increased frameshift indel burden compared with ncHRD (p<0.01) and HRP (p<0.001) tumors (**Figure 2E i**). Consistent with this enrichment, HRD tumors also demonstrated a higher number of predicted major histocompatibility (MHC) class I-binding neoantigens, despite relatively modest differences in overall TMB across cohorts (**Figure 2E ii**).

Across the combined discovery (iPC) and validation (POLAR) cohorts, predicted neoantigen burden demonstrated a strong positive correlation with MDF in HRD PC (Pearson R=0.72, p=0.00078). In contrast, weaker or non-significant associations were observed in ncHRD and HRP tumor (**Figure 2F**), suggesting that MDF represents an important source of frameshift indel-derived high-quality neoantigens specifically in HRD PC. To determine whether this association was simply driven by overall mutation burden, we adjusted neoantigen burden for TMB. MDF remained positively associated with residual neoantigen burden after accounting for TMB (**Figure S5C**), supporting that POLQ-mediated deletion scars preferentially generate immunogenic frameshift-derived neoantigens beyond the effects of mutation quantity alone.

A conceptual model integrating these observations is presented in **Figure 2B**. In HRD tumors, unresolved double-strand breaks may be preferentially repaired through error-prone MMEJ, generating short deletions that frequently produce frameshift indels and predicted neoepitopes capable of promoting baseline immune engagement. Notably, the magnitude of neoantigen enrichment exceeds what would be expected from the modest differences in overall TMB, suggesting that the qualitative nature of MMEJ-associated frameshift indels rather than TMB drives neoantigen generation in HRD tumors.

### Neoantigen burden in HRD PC aligns with HRD and MDF metrics, independent of plastic subtypes

To contextualize MMEJ-driven immunogenicity within established genomic frameworks, we evaluated neoantigen burden across multiple HRD-related stratifications. Neoantigen burden was significantly associated with HRD gene zygosity status (biallelic vs monoallelic; p<0.01) and MDF high versus low scores (p<0.001) (**Figure S5D**). However, there was no association with IMPACT-HRD score or *BRCA* genotype (*BRCA2* vs *BRCA1* vs *PALB2)*, suggesting that variation in neoantigen burden within HRD PC is better captured by MMEJ repair activity quantified by MDF.

By contrast, established PC transcriptional subtypes defined by malignant cell plasticity using the PurIST classification were largely independent of genomic HRD features. Distribution of classical and basal-like tumors did not differ significantly by HRD status, IMPACT-HRD score or neoantigen burden (**Figure S6A)**. Although classical tumors exhibited a trend toward improved survival prognosis compared with basal-like tumors (**Figure S6C-D**), this occurred despite comparable IMPACT-HRD scores and neoantigen burdens. These observations indicate that transcriptional plasticity and genomic instability represent distinct biological axes in pancreatic cancer.

Notably, no tumors with adenosquamous or squamous histology were included in this cohort. Therefore, basal-like transcriptional programs observed here reflect plastic malignant cell states within conventional pancreatic acinar cell carcinoma and adenocarcinoma rather than distinct histologic variants^28^. Together, these analyses support MDF as a genomic marker of DNA repair-associated immunogenicity in HRD PC that is independent of both canonical HRD metrics and transcriptional subtype classifications.

### MDF-derived neoantigens shape the immune landscape of HRD pancreatic cancer

To define how POLQ-mediated genomic scarring and the resultant neoantigen burden shape the tumor immune microenvironment, we analyzed immune cell nuclei profiled by single nucleus RNA sequencing (snRNA-seq) from HRD and HRP PCs in the iPC cohort (**Figure 3; Figure S7**). Across 34,677 immune cell nuclei, we identified major immune compartments including CD4⁺ T cells, CD8⁺ T cells, B cells, macrophages, NKT cells and regulatory T cells, with further resolution of functional states spanning naïve, cytotoxic, effector, memory and exhausted phenotypes (**Figure 3A–B**). Among the 23 tumors profiled by snRNA-seq, a subset of eight tumors had matched WES enabling neoantigen annotation and MDF calculation and these samples were used for integrative analyses linking genomic features with immune cell states.

**Figure 3.**
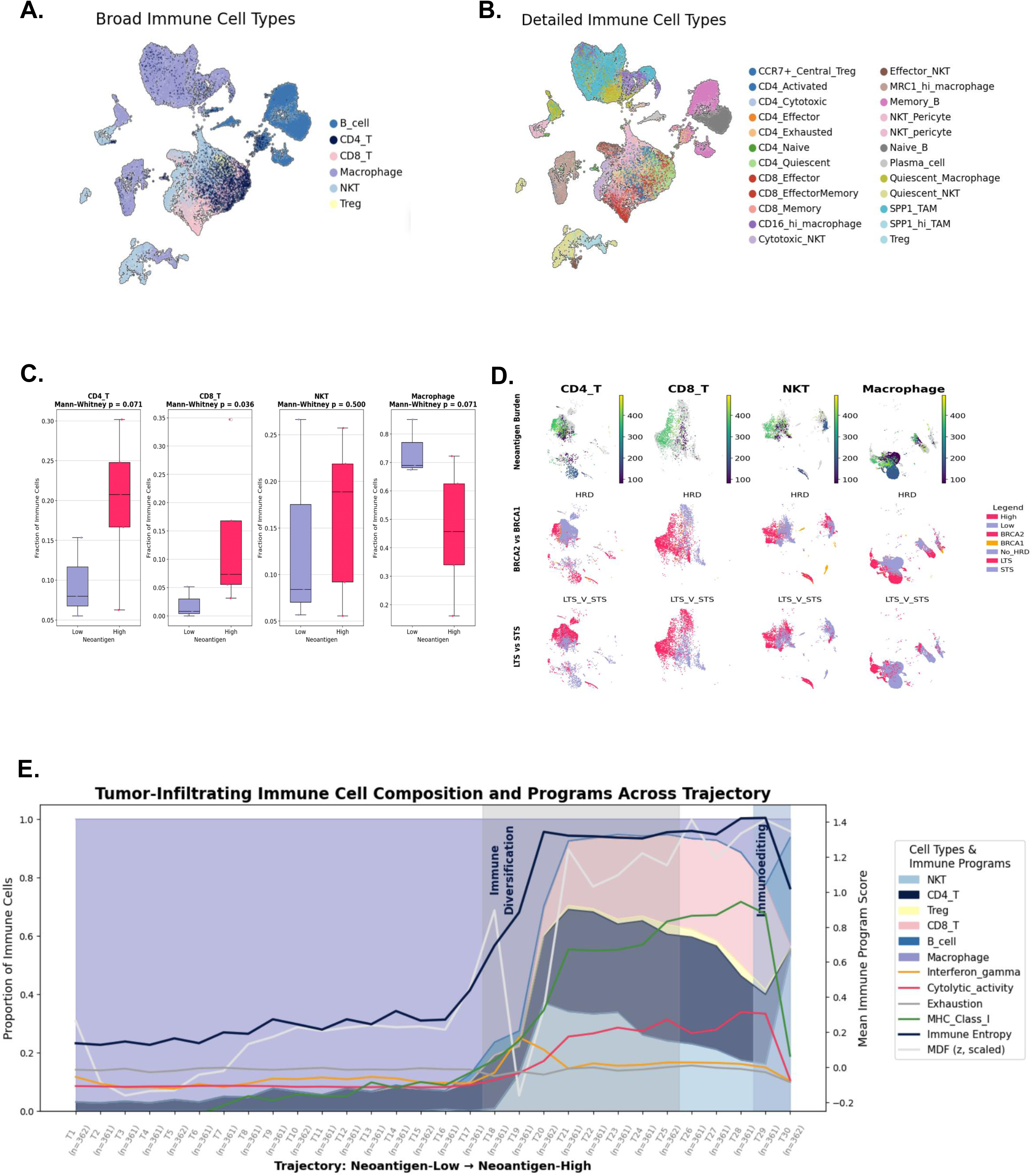
MDF-derived neoantigens shapes immune landscape of HRD pancreatic cancer. **(A–B)** UMAP of immune cells (N = 52,309) across single-nucleus RNA-seq (snRNA-seq) from HRD and HRP pancreatic cancers, colored by (A) broad and (B) detailed immune compartments. **(C)** Boxplots reveal that neoantigen-high tumors are enriched for CD8⁺ T cells (p=0.036), CD4+T cells (p=0.071), and exhibit decreased macrophage infiltration (p=0.071), supporting an immunogenic tumor microenvironment (TME), explaining the inverse relationship of higher immune infiltrates in low neoantigen tumors was mostly from the macrophages. **(D)** UMAP overlays of immune cell subsets stratified by: **Top row:** Neoantigen burden (high vs. low), **Middle row:** BRCA2 vs BRCA1 mutations within HRD tumors, **Bottom row:** Long-term survivors (LTS) vs short-term survivors (STS). **(E)** Multi-layered immune cell entropy and trajectory analysis plot showing immune cell proportions (stacked area) at each trajectory point within the increasing order of neoantigen trajectory. The immune program scores (line overlay) across immune trajectory bins ordered by neoantigen burden. The transition from macrophage-dominant to T cell–rich microenvironments coincide with rising MHC class I/II, IFN-γ, and cytolytic activity, followed by a decline in these programs and immune entropy, consistent with immunoediting dynamics. **Abbreviations:** HRD, homologous recombination–deficient; HRP, homologous recombination–proficient; NKT, natural killer T cell; TAM, tumor-associated macrophage; TME, tumor microenvironment; LTS, long-term survivor; STS, short-term survivor; snRNA-seq, single-nucleus RNA sequencing.

Tumors with high neoantigen burden exhibited enrichment of CD8⁺ (p=0.036) and CD4⁺ T cells (p=0.071) and relative depletion of tumor-associated macrophages (TAMs) (p=.071), consistent with an immunogenic tumor microenvironment (**Figure 3C**). Projection of neoantigen burden, *BRCA1/2* genotype and survival status onto immune UMAP embeddings demonstrated that neoantigen-high tumors preferentially occupied immune-activated regions enriched for CD8⁺ T and NKT cells, whereas neoantigen-low tumors were dominated by macrophage-rich states (**Figure 3D**). Within HRD tumors, *BRCA2*-mutant cancers and long-term survivors (overall survival > 18 months) were disproportionately represented in immune-activated regions, linking genomic context, immune state and clinical outcome.

To integrate immune composition with functional programs, immune cells from tumors with matched neoantigen data (N=8) were ordered along a neoantigen-associated immune gradient and summarized cells across 30 ordered trajectory bins rather than individual tumors (**Figure 3E**). This approach captures progressive immune-state changes across the pooled immune compartment while preserving each tumor’s contribution according to its cellular composition. Along this trajectory, we observed a transition from macrophage-predominant states toward a more immune-diverse state enriched for NKT, CD4⁺ T, CD8⁺ T and B cells. This was accompanied by increased antigen presentation, interferon-γ signaling and cytolytic activity. At later trajectory stages, these programs declined sharply, consistent with immunoediting following sustained immune pressure. Mean tumor-level MDF increased in parallel with immune activation and diversification before plateauing, supporting a model in which POLQ-mediated repair scars and derived neoantigens promote immune recruitment, activation and subsequent immune sculpting over time.

### Neoantigen burden polarizes CD8⁺ T cell state

To determine how neoantigen burden shapes adaptive immune responses in HRD PC, we focused on tumor-infiltrating CD8⁺ T cells identified by snRNA-seq in the iPC cohort **(Figure 4**). These analyses were performed on pretreatment biopsy specimens collected prior to standard treatment exposure^24^, thereby capturing the baseline immune landscape without therapeutic immune modulation. Biopsy timing and site distribution are summarized in **Table S2**.

**Figure 4.**
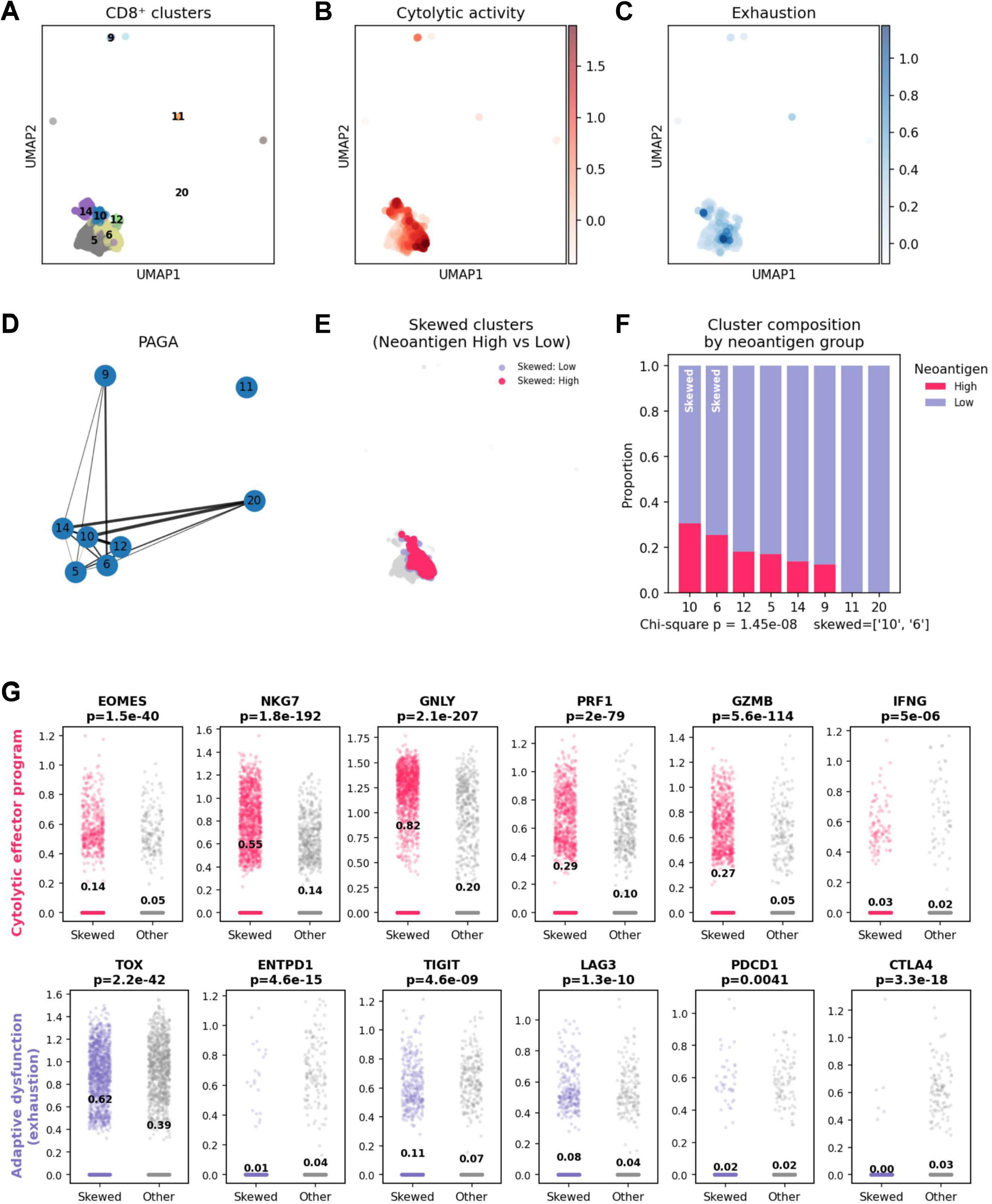
Neoantigen burden is associated with cytotoxic and dysfunctional polarization of tumor infiltrating CD8+ T cells in HRD pancreatic cancer. (A) UMAP of tumor infiltrating CD8⁺ T cells colored by Leiden clusters demonstrates transcriptionally distinct CD8⁺ T cell states. (B) Cytolytic activity module score projected on the UMAP highlights enrichment of effector programs within a subset of CD8⁺ T cells. (C) Exhaustion module score projected on the UMAP demonstrates heterogeneity in adaptive dysfunction programs across CD8⁺ T cell states. (D) PAGA graph summarizes inferred connectivity between Leiden clusters, consistent with divergent differentiation trajectories within the CD8⁺ T cell compartment. (E) Cells from the neoantigen enriched CD8⁺ T cell clusters (Leiden clusters 10 and 6) are shown on the UMAP and colored by tumor neoantigen group (high vs low). (F) Stacked bar plot quantifies cluster level composition by neoantigen group. Skewed clusters (clusters 10 and 6) are relatively enriched in neoantigen high tumors compared with other clusters (Chi-square P = 1.45 × 10⁻⁸). (G) Expression of cytolytic effector genes (top row) and adaptive dysfunction or exhaustion associated genes (bottom row) in skewed versus other CD8⁺ T cells. Points represent single cells and values denote group means with two-sided Mann Whitney P values shown

Unsupervised clustering of CD8⁺ T cells identified transcriptionally distinct clusters that, based on cytolytic and exhaustion gene programs, corresponded to naïve-like, cytotoxic effector and dysfunctional or exhausted phenotypes (**Figure 4A-C**). Projection of functional gene programs onto the CD8⁺ T cell UMAP revealed marked polarization of cytolytic activity and adaptive dysfunction (**Figure 4B–C**). Cytolytic activity, defined by coordinated expression of *PRF1, GZMB, GNLY, NKG7* and *EOMES,* localized to a discrete subset of CD8⁺ T cells, whereas exhaustion-associated programs including *TOX, LAG3, TIGIT, PDCD1* and *CTLA4* exhibited overlapping but non-identical distributions. These connections suggest that cytotoxic and dysfunctional CD8⁺ T cell states arise along a continuous differentiation trajectory rather than representing separate terminal populations.

Graph-based abstraction of CD8⁺ T cell state transitions using PAGA revealed structured connectivity among clusters, consistent with a continuum of antigen-associated differentiation within the CD8⁺ compartment **(Figure 4D**). Cytolytic effector–enriched clusters were directly connected to clusters exhibiting adaptive dysfunction, indicating that functionally constrained or exhausted states emerge along a shared differentiation axis rather than as abrupt terminal fates.

Overlay of tumor neoantigen burden onto the CD8⁺ T cell UMAP demonstrated preferential enrichment of cells from neoantigen-high tumors within a subset of clusters, most prominently Leiden clusters 10 and 6 **(Figure 4E**). These clusters corresponded to antigen-primed cytotoxic/dysfunctional CD8^+^T cell states characterized by elevated cytolytic effector genes together with adaptive dysfunction markers. Quantitative analysis confirmed that these skewed clusters were significantly enriched in neoantigen-high tumors compared with other CD8⁺ T cell states (Chi-square p=1.45 × 10⁻⁸; **Figure 4F**), and CD8⁺ T cells within these clusters expressed significantly higher *PRF1, GZMB, GNLY, NKG7, TOX, TIGIT and LAG3* relative to other CD8⁺ T cells (**Figure 4G**).

Together, these data demonstrate that higher neoantigen burden in HRD pancreatic cancer is associated with selective polarization of CD8⁺ T cells toward antigen-primed effector–dysfunctional states present at baseline, consistent with pre-existing tumor immunogenicity rather than *de novo* treatment-induced immune activation.

### Neoantigen burden reshapes macrophage programs and immune synapse architecture

Given the reciprocal relationship between T cell activation and myeloid cell states, we next examined macrophage transcriptional programs as a function of tumor neoantigen burden (**Figure 5A-B; Figure S8)**. Conventional dendritic cells were rare in this snRNA-seq dataset, and to avoid conflating macrophage states with antigen-presenting cell (APC)-like populations, we excluded the MHC II-high DC-like macrophage subset from this macrophage-focused clustering analysis.

**Figure 5.**
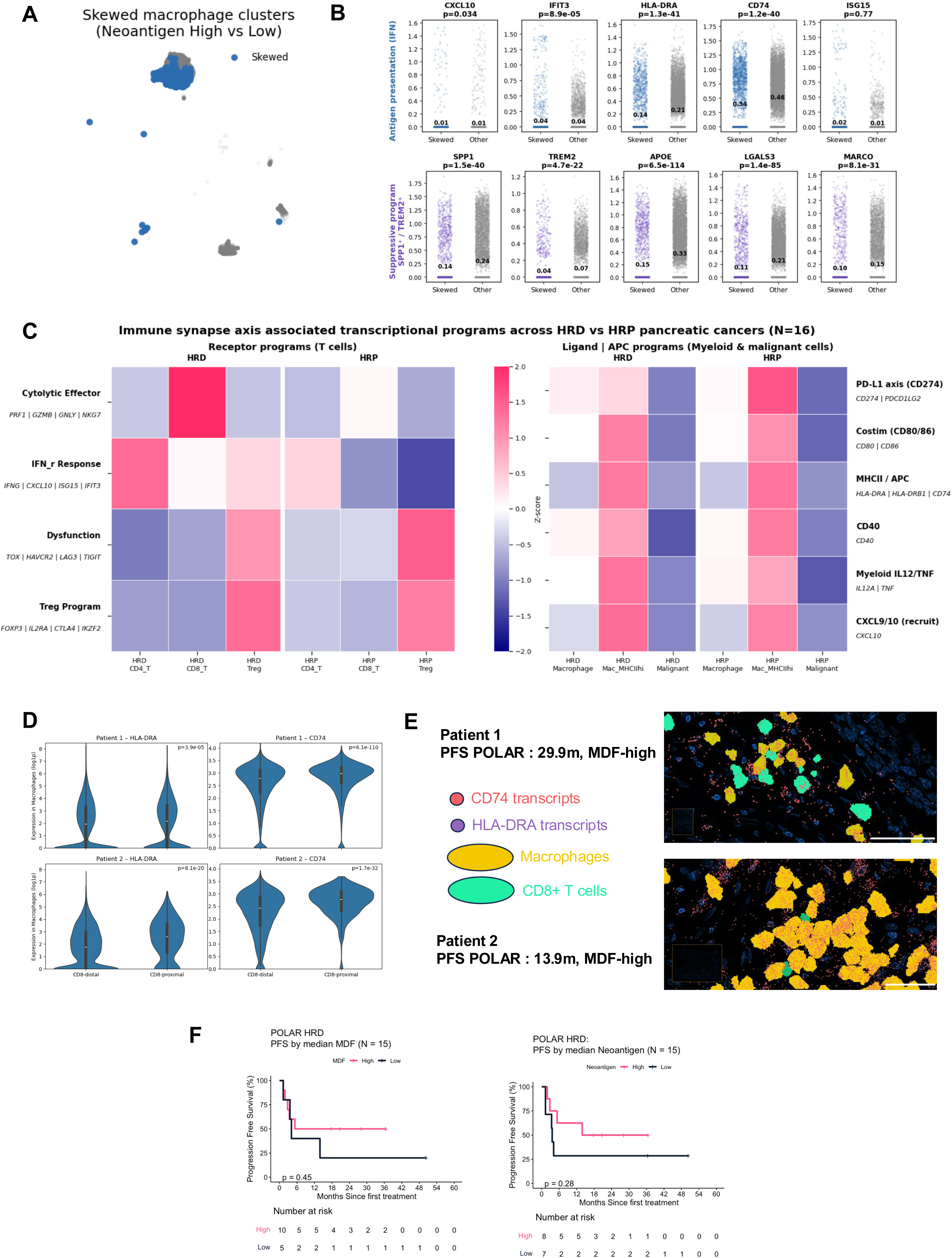
Intrinsic immunogenic programs interface with immune synapse architecture and associate with clinical outcome. **(A)** UMAP highlighting macrophage clusters selectively enriched in tumors with high neoantigen burden (“skewed” clusters; blue) compared with other macrophage populations (gray). **(B)** Differential expression of antigen presentation–associated genes (top row) and immunosuppressive macrophage programs (bottom row) comparing skewed versus other macrophage clusters. Each point represents a single cell; bars indicate mean expression per group. Two-sided Mann–Whitney U tests are shown. MHC II–high dendritic cell–like macrophages were excluded from this macrophage-focused analysis. **(C)** Heatmap summarizing immune synapse–associated transcriptional programs across homologous recombination–deficient (HRD; N = 10) and homologous recombination–proficient (HRP; N = 6) pancreatic cancers within the snRNA-seq cohort (total N = 16 samples only at baseline with germline HRD annotation available). Left: receptor-side programs in T cells. Right: ligand and antigen-presenting programs in myeloid and malignant cells. Values represent z-scored, sample-aggregated program activity. **(D)** Violin plots showing HLA-DRA and CD74 expression (log1p-normalized) in macrophages stratified by proximity to CD8⁺ T cells (≤20 µm vs >20 µm) in two baseline primary tumors from POLAR long-term responders. P-values were computed using two-sided Mann–Whitney U tests. **(E)** Representative spatial transcriptomic images from two treatment-naïve primary pancreatic tumors obtained at baseline from metastatic patients enrolled in the POLAR trial. Each panel corresponds to one individual patient. Macrophages are shown in yellow and CD8⁺ T cells in cyan. Transcripts of **HLA-DRA** (purple) and **CD74** (red), key components of the MHC class II antigen presentation machinery, are overlaid at single-molecule resolution. In both patients, CD8⁺ T cells are frequently found in close spatial proximity to macrophages expressing HLA-DRA and CD74, consistent with localized antigen-presenting niches within the tumor microenvironment. Scale bars, 50 µm. **(F)** Progression-free survival (PFS) in HRD pancreatic cancer (POLAR cohort; N = 15) stratified by median MDF (top) and median neoantigen burden (bottom). Neoantigen burden demonstrated a trend toward improved PFS, with MDF showing a concordant but weaker association. Overall survival analyses were exploratory due to limited event maturity. **See also Figure S8.**

Within the remaining macrophage population, unsupervised clustering resolved transcriptionally distinct subsets dominated by tissue remodeling and suppressive programs, including *SPP1^hi^TREM2^hi^*state, opposed to classical interferon-activated or antigen-presenting macrophage states **(Figure 5A**). Consistent with this filtering, interferon-related and antigen presentation genes were not increased in macrophages from neoantigen-high tumors relative to other macrophage subsets in this analysis (**Figure 5B**). Instead, macrophage populations enriched in neoantigen-low tumors exhibited higher expression of *SPP1, TREM2, APOE, LGALS3* and *MARCO*, consistent with an immunosuppressive and tissue-remodeling macrophage programs (**Figure 5B; Figure S8B**).

Differences in macrophage cluster composition by neoantigen group remained highly significant (Chi square p=4.19 × 10⁻³⁵; **Figure S8F**), indicating that neoantigen burden is associated with structured remodeling of macrophage states. Because APC-like macrophages were excluded from this clustering analysis, these results primarily reflect changes in suppressive macrophage programs rather than antigen-presenting macrophage states. APC-like macrophages therefore represent a distinct antigen-presenting niche within the tumor microenvironment and were examined separately in downstream immune synapse analyses.

### Immune synapse programs align with neoantigen-driven immune polarization and clinical outcome

To integrate lymphoid and myeloid programs within a unified framework, we evaluated immune synapse–associated transcriptional programs across baseline HRD and HRP PC samples with germline HRD annotation available (N=16), using sample-aggregated gene module scoring (**Figure S8G**). Receptor-side programs (**Figure 5C**, left) in T cells–including cytolytic effector, interferon response, dysfunction and Treg-associated modules–were selectively elevated in HRD tumors within both CD8⁺ T and CD4⁺ T cell compartments, consistent with the polarized T cell states described above.

In parallel, ligand and antigen-presenting programs (**Figure 5C**, right) in macrophages and malignant cells–including PD-L1 (CD274), costimulatory molecules (CD80/CD86), MHC II, CD40 and CXCL9/10–were coordinately upregulated in HRD tumors relative to HRP tumors. The coordinated enrichment of receptor-side T cell programs and ligand/APC programs across compartments suggests structured immune communication, more than simple immune cell infiltrations alone.

To directly assess whether MDF- and neoantigen-high HRD PCs exhibit spatial immune organization consistent with productive antigen presentation, we performed spatial transcriptomic profiling on baseline, treatment-naïve primary tumor biopsies from two POLAR participants with prolonged clinical benefit (progression-free survival of 13.9 for patient 1 and 29.9 months for patient 2) on the pembrolizumab and olaparib maintenance. Both tumors had high MDF and neoantigen burden. Using single-cell–resolved spatial coordinates, we quantified the proximity of CD8⁺ T cells to APC–like macrophages, defined by macrophages expressing both HLA-DRA (p=3.9 x 10^-5^ and p=8.1 x 10^-20^) and CD74 (p=6.1 x 10^-110^ and p=1.7 x 10^-32^). In a tumor from patient 2, CD8⁺ T cells were significantly enriched within 20 µm of APC-like macrophages relative to a spatial permutation null model that preserved tissue geometry and cell density. This is consistent with structured immune synapse niche. In contrast, the second tumor did not exhibit significant CD8⁺ T cell enrichment.

Importantly, even in the absence of overt spatial enrichment of CD8⁺ T cells, macrophages proximal to CD8⁺ T cells in both tumors displayed markedly higher expression of antigen-presenting markers. In both tumors, CD8⁺ T-proximal macrophages (<20 µm from CD8⁺ T cells) displayed significantly higher expression of HLA-DRA and CD74 compared to CD8⁺ T-distant macrophages (> 20 µm; both p<10⁻¹¹), with a markedly larger effect size in the tumor exhibiting spatial CD8⁺ T–APC enrichment (**Figure 5D**). These findings suggest that CD8⁺ T cell proximity is associated with localized reinforcement of antigen-presenting programs with macrophages, supporting the presence of functional immune synapse architecture.

Together, these analyses reveal two complementary modes of immune organization in MDF- and neoantigen-high HRD pancreatic cancer: one characterized by structured spatial enrichment of CD8⁺ T cells around APC-like macrophages, and another marked by diffuse but locally reinforced antigen presentation. Both configurations are consistent with functional immune synapse architecture and provide a spatial correlate to the neoantigen-driven immune activation observed in these long-term responders.

Finally, within the HRD POLAR cohort (N = 15), higher neoantigen burden was associated with prolonged progression-free survival, whereas MDF demonstrated a directionally concordant but weaker association (**Figure 5E**). Restricted mean survival time (RMST) analysis over 24 months demonstrated longer progression free survival in neoantigen high tumors compared with neoantigen low tumors (difference 6.2 months), with MDF high tumors showing a smaller concordant difference (4.1 months). Given the small cohort size and limited event maturity, these differences did not reach statistical significance and overall survival analyses were considered exploratory. Together, these data link POLQ-mediated genomic scarring and neoantigen burden to coordinated immune synapse engagement and clinical outcome in HRD pancreatic cancer.

## DISCUSSION

Pancreatic cancer remains one of the most immune-refractory solid tumors, with immune checkpoint blockade (ICB) demonstrating limited efficacy in unselected populations^3,29–32^. This resistance reflects a convergence of barriers, including low baseline neoantigen burden, dominant myeloid suppression, and stromal exclusion, which collectively constrain productive T-cell priming and engagement^33,34^. Within this landscape, DNA damage repair (DDR)–deficient subgroups, including mismatch repair deficiency (dMMR) and homologous recombination deficiency (HRD), are uncommon but mechanistically-poised contexts in which tumor-intrinsic immunogenic signals may emerge^7,35,36^. However, the specific genomic features within DDR-deficient tumors that generate sustained and functionally relevant tumor immunogenicity remain incompletely defined.

Here, we demonstrate that genomic repair scars generated by POLQ-mediated microhomology-mediated end joining (MMEJ) repair scars, quantified as the MMEJ Deletion Footprint (MDF), provide a mechanistic bridge linking HRD to neoantigen quality, coordinated immune activation and clinically meaningful baseline immune engagement in pancreatic cancer (PC).

By integrating exome-derived mutational features, predicted neoantigen properties, bulk transcriptional programs, and single-nucleus resolution of malignant and immune compartments, we define a coherent immunogenic axis in HRD pancreatic cancer. We identify microhomology-mediated deletion scars, quantified as the MMEJ deletion footprint (MDF), as a key genomic consequence of double-strand break repair through an error-prone alternative pathway, which generates frameshift indel neoantigens and malignant cell-intrinsic programs of interferon signaling and antigen presentation. Importantly, these cell-intrinsic programs are mirrored by coordinated remodeling of the tumor microenvironment, including selective polarization of CD8⁺ T cells toward cytolytic, antigen-primed states and alignment with macrophage programs capable of supporting antigen presentation and immune synapse engagement. Immune activation in this setting is not attributable to any single cellular compartment but instead emerges from the coordinated alignment of genomic scarring, antigenic output, innate nucleic acid sensing and adaptive immunity.

A central conceptual insight is that not all mutations contribute equally to immune recognition, with frameshift indels being disproportionately capable of generating high-affinity non-self epitopes, even in tumors with modest overall mutational burden^11,12,19,26,27,37^. In HRD pancreatic cancer, clonal driver mutations such as *KRAS, TP53* and *CDKN2A* still dominate the genomic landscape, whereas repair-associated passenger mutations accumulate over time. Among these, MMEJ-derived frameshift indels provide a potent substrate for neoantigen generation^38,39^. Our data support a model in which neoantigen quality, rather than mutation count alone, aligns with tumor-intrinsic antigen presentation and interferon-γ–associated programs and extends downstream to productive CD8⁺ T cell activation. This framework helps explain why a subset of HRD tumors exhibits functional immune engagement despite the broader resistance of pancreatic cancer to immunotherapy^19,40,41^.

Beyond adaptive immune recognition, the immunogenic features observed in MDF-high tumors are consistent with tumor-intrinsic stress responses associated with interferon signaling and antigen presentation pathways. Such responses are frequently linked to nucleic acid sending triggered by genotoxic stress in the setting of DNA repair deficiency^42–44^. These sensing pathways converge on TBK1–IRF3 signaling and induce transcriptional programs that reinforce antigen presentation and adaptive immune engagement^45,46^. Consistent with this framework, MDF-high tumors showed transcriptional evidence of interferon-associated immune activation alongside enhanced antigen presentation program. These observations suggest that HRD-associated genomic scarring may promote immunogenicity not only through neoantigen generation but also through tumor-intrinsic immune signaling linked to genotoxic stress.

Our analyses further highlight antigen-presenting macrophage states as a critical execution layer, linking tumor intrinsic immunogenicity to adaptive immune engagement. Neoantigen-enriched tumors align with macrophage programs consistent with antigen presentation and inflammatory chemokine output, whereas neoantigen-low tumors remain dominated by suppressive SPP1^hi^ and TREM2^hi^ states^47,48^. APC-like macrophages form a transcriptionally distinct program rather than a continuum of suppressive TAM states; accordingly, they were excluded from macrophage trajectory analyses focused on TAM polarization but incorporated in downstream analyses examining immune synapse signaling. This distinction is clinically relevant, as ineffective antigen presentation and suppressive myeloid signaling can constrain durable cytolytic control even in the presence of infiltrating T cells. The coordinated alignment of T cell receptor programs with ligand and costimulatory programs in macrophages and malignant cells support an immune synapse axis in which PD1-PDL1 interactions, costimulation and antigen presentation programs co-occur in specific cellular contexts. These findings suggest that productive immunity in HRD PC depends not only on T cell presence, but on whether myeloid cells occupy APC-like states capable of sustaining effective immune synapse engagement. In contrast to the continuous differentiation trajectory observed within the CD8^+^T-cell compartment, APC-like macrophages appear to occupy a discrete functional niche.

Importantly, immune engagement in this context is defined not solely by immune cell abundance, but by the spatial organization of immune interactions consistent with an immune synapse architecture. We propose a model in which HRD establishes the genomic substrate, MDF and neoantigen features drive tumor intrinsic activation and antigen presentation, and macrophage state enables effective CD8⁺ T cells and HLA-DR+ macrophages, together with enrichment of checkpoint and costimulatory programs, provides a measurable architecture that may be more informative than bulk sequencing metrics. This is consistent with growing evidence across solid tumors that neighborhood interactions best capture functional antitumor immunity^49,50^. Notably, such immune synapse architecture can be assessed using spatial transcriptomics or multiplex imaging platforms, supporting translational feasibility.

Although MMEJ is classically associated with HR deficiency^22,51,52^, we detected low-level MDF signal in a minority of HR-proficient tumors; however, its coupling to neoantigen burden was markedly weaker than in HRD tumors. These events likely reflect episodic engagement of error-prone repair under conditions of replication stress rather than a dominant mutational process^22,51,53^. In this setting, infrequent MMEJ events may still generate frameshift deletions, but their aggregate contribution appears limited. These observations reinforce the interpretation that MDF is most informative when contextualized within HRD and underscore the need for larger datasets with orthogonal measures of replication stress and repair pathway engagement to define its broader applicability.

Although transcriptional subtype plasticity is an important determinant of pancreatic cancer biology, immune programs did not segregate by subtype in our cohort^38,54^. Classical and basal-like features were not predictive of HRD status or MMEJ deletion footprint. In contrast, the MDF-associated immunogenic axis – spanning neoantigen burden, interferon signaling and antigen presentation – was evident across heterogeneous transcriptional states. These findings argue that the immune associations observed here reflect DDR-linked mutational processes rather than plastic subtype-specific inflammatory or stromal programs.

These findings also raise the possibility that therapeutic DNA damage may amplify the immunogenic consequences of error-prone repair in HRD tumors. Platinum chemotherapy induces extensive double-strand DNA breaks that must be resolved through alternative repair mechanisms when HRD. In this setting, MMEJ represents a key compensatory repair mechanism. Consistent with this, recent analyses of niraparib-treated HRD pancreatic cancers have shown that *BRCA2* reversion mutations frequently arise through MMEJ-mediated repair events that restore the open reading frame of the mutated gene^23,55^. Although such reversion mutations represent a mechanism of niraparib resistance, they highlight the MMEJ activity in HRD PC under therapeutic pressure. We speculate that the same repair processes that occasionally generate reversion mutations may more commonly produce deletion scars and frameshift events capable of generating high quality neoantigens. In this framework, induction platinum-based chemotherapy can generate neoantigens^56^, potentially creating an exploitable immunogenicity. This concept is relevant to post-platinum maintenance strategies like POLAR trial^19^, in which patients receiving induction platinum-based chemotherapy subsequently switched to olaparib and pembrolizuamb maintenance during the period of chemically and molecularly minimal disease.

This study has limitations, including modest cohort sizes for select analyses, reliance on exome-based inference for neoantigen prediction and incomplete availability of matched spatial material across all cases. Nevertheless, exome-derived features are directly relevant to neoantigen discovery and broadly applicable in clinical settings. Within these constraints, our analyses consistently link POLQ-associated repair scarring to functional immune engagement in HRD pancreatic cancer. Prospective studies will be required to establish the predictive utility of MDF and immune synapse organization for immunotherapy response; however, these data provide a strong biological rationale for combination strategies that augment antigen presentation alongside immune checkpoint blockade.

## CONCLUSIONS

Together, these findings identify POLQ-mediated MMEJ repair scars as a mechanistic axis linking homologous recombination deficiency to neoantigen quality, immune synapse organization and functional antitumor immunity in pancreatic cancer. HRD establishes the conditions that favor error-prone DNA repair, generating frameshift neoantigens capable of initiating coordinated innate and adaptive immune responses. By connecting DNA repair biology to tumor immunogenicity, this framework provides a biological foundation for precision immunotherapy strategies in HRD pancreatic cancer and potentially other homologous recombination-deficient malignancies.

## METHODS

### Study cohorts and clinical annotation

We analyzed PDAC specimens from (i) a multi-omic discovery cohort (iPC) with matched tumor whole-exome sequencing (WES) and single-nucleus RNA sequencing (snRNA-seq) and available clinical annotation, and (ii) an independent HRD-enriched clinical cohort from the POLAR study with tumor WES and longitudinal clinical outcomes. HRD classification required pathogenic or likely pathogenic variants in core homologous recombination genes (*BRCA1*, *BRCA2, PALB2*) or selected HRR genes. dMMR/MSI-H cases were identified by standard clinical or sequencing-based criteria. Clinical endpoints included [PFS/OS] as defined in the study protocols. All participants provided informed consent under Institutional Review Board–approved protocols at Memorial Sloan Kettering Cancer Center.

### Whole-exome sequencing processing and variant calling

WES data were processed using a standardized pipeline. Reads were aligned to the human reference genome (GRCh38) using BWA-MEM. Duplicate marking and base quality recalibration were performed using GATK best practices. Somatic single nucleotide variants and indels were called using [Mutect2 / Strelka2] with matched normal samples when available. Variants were filtered using panel-of-normals and artifact filters and annotated using [VEP/Oncotator]. For samples without matched normals, additional filtering was applied using population databases and artifact priors.

### Definition of the Microhomology Mediated End Joining Deletion Footprint

Somatic deletion lengths were calculated from whole-exome sequencing data by comparing the reference and alternate allele lengths for each deletion event. For each tumor, deletions were grouped by size, including 1–5 bp, 6–13 bp, 14–20 bp and 21–50 bp bins. These deletion size ranges correspond to the characteristic microhomology-mediated deletion spectrum previously associated with MMEJ repair in human tumors. The MMEJ Deletion Footprint (MDF) was defined as the summed count of deletions in the 6–13 bp and 14–20 bp bins, corresponding to the characteristic deletion size range associated with MMEJ repair. Raw MDF counts were standardized across samples using z-score normalization and shifted to positive values for visualization. MDF quartiles were defined using the standardized MDF values.

### Neoantigen prediction and frameshift neoantigen prioritization

HLA class I alleles were inferred from WES using POLYSOLVER. Candidate neoantigens were generated from expressed coding variants, including frameshift indels. Peptide binding affinity to HLA class I was predicted using NetMHCpan (version 4.0). Neoantigens were prioritized based on predicted binding affinity and rank, peptide processing heuristics, and clonality surrogates when available.

### snRNA-seq processing, clustering, and cell state annotation

snRNA-seq data were processed using Cell Ranger followed by downstream analysis in Seurat/Scanpy. Nuclei were filtered by UMI counts, gene counts, and mitochondrial read fraction. Data were normalized and integrated across samples using Harmony. Dimensionality reduction was performed using PCA followed by UMAP. Clustering was performed using Leiden/Louvain. Major cell compartments (malignant, immune, stromal, endothelial) were annotated using canonical marker expression and module scoring. Malignant cell identity was further supported using inferred copy number profiles derived from transcriptional data.

### Immune programs, macrophage states, and CD8 functional trajectories

Gene expression programs including antigen presentation (MHC I/II), interferon response, cytotoxicity, exhaustion, and innate sensing signatures were quantified using module scoring with curated gene sets. Macrophage states were defined by expression of antigen presentation genes and suppressive programs including SPP1 and TREM2-associated signatures. CD8 T cell states were assessed using established markers and module scores, and where trajectory analyses were performed, pseudotime was inferred using [Monocle3 / Slingshot / Palantir], with root states defined as [naïve/low cytotoxicity] CD8 populations.

### Spatial profiling and immune synapse architecture

Spatial transcriptomic analyses were performed on baseline, treatment-naïve primary tumor sections from two metastatic patients enrolled in the POLAR trial, using the 10x Genomics Xenium platform. Spatially resolved single-cell coordinates and gene expression matrices were used for all analyses. Cell types were annotated based on curated cell-type labels. APC-like macrophages were stringently defined as macrophages co-expressing HLA-DRA and CD74, using log1p-normalized expression values from the primary expression matrix (expression > 0 for both genes).

To assess spatial proximity between CD8⁺ T cells and APC-like macrophages, we quantified the fraction of CD8⁺ T cells located within a 20 µm radius of at least one APC-like macrophage using Euclidean distances computed from spatial coordinates expressed in micrometers. To evaluate whether observed proximity exceeded that expected by chance, we implemented a permutation-based spatial null model in which APC-like labels were randomly reassigned among macrophages while preserving overall cell positions, tissue geometry, and cell density. For each sample, 500 permutations were performed, and empirical p-values were calculated as the proportion of permuted values greater than or equal to the observed statistic.

To evaluate proximity-dependent activation of antigen presentation programs, macrophages were stratified into CD8-proximal (≤20 µm to the nearest CD8⁺ T cell) and CD8-distal (>20 µm) populations. Expression levels of HLA-DRA and CD74 were compared between proximal and distal macrophages within each sample using two-sided Mann–Whitney U tests. All spatial analyses were performed independently for each sample to avoid confounding by inter-patient variability. Statistical analyses were conducted using Python (SciPy), and data visualization was performed using Matplotlib and Seaborn.

### Statistical analysis

Group comparisons were performed using nonparametric tests, including Mann–Whitney U tests for continuous variables and Fisher’s exact tests for categorical variables where appropriate. Correlations between continuous variables were evaluated using Spearman rank correlation. Survival analyses were performed using Kaplan–Meier estimation with log-rank testing and Cox proportional hazards models. Restricted mean survival time (RMST) analyses were performed with truncation time τ = 24 months using the survRM2 package in R.

Unless otherwise indicated, statistical tests were two-sided and P<0.05 was considered statistically significant. Statistical analyses were conducted using R (version 4.3.2) and Python (version 3.10).

## Data and code availability

Sequencing data will be deposited at dbGaP under accession [x], subject to controlled access where required. Processed data and analysis code will be made available at https://github.com/WilliamPolar/MDF-immunogenicity-PC.

## ACKNOWLEDGEMENTS

We are deeply grateful to the patients and families who participated in these studies and whose generosity made this work possible. We thank the clinical research staff, study coordinators, nurses and laboratory personnel at Memorial Sloan Kettering Cancer Center for their dedication to patient care and biospecimen collection. We thank the members of the gastrointestinal oncology and translational research teams for their contributions to patient enrollment, sample processing, and data generation.

We are particularly grateful to the Katz family for their philanthropic support of pancreatic cancer research. This work was supported in part by the National Institutes of Health/National Cancer Institute Pancreatic Cancer SPORE (P50 CA257881) and by institutional and philanthropic support from the David M. Rubenstein Center for Pancreatic Cancer Research at Memorial Sloan Kettering Cancer Center. We acknowledge the Integrated Genomics Operation Core at Memorial Sloan Kettering Cancer Center for sequencing services, supported by the NCI Cancer Center Support Grant (P30 CA008748)

## TABLE LEGENDS

**Table S1.**
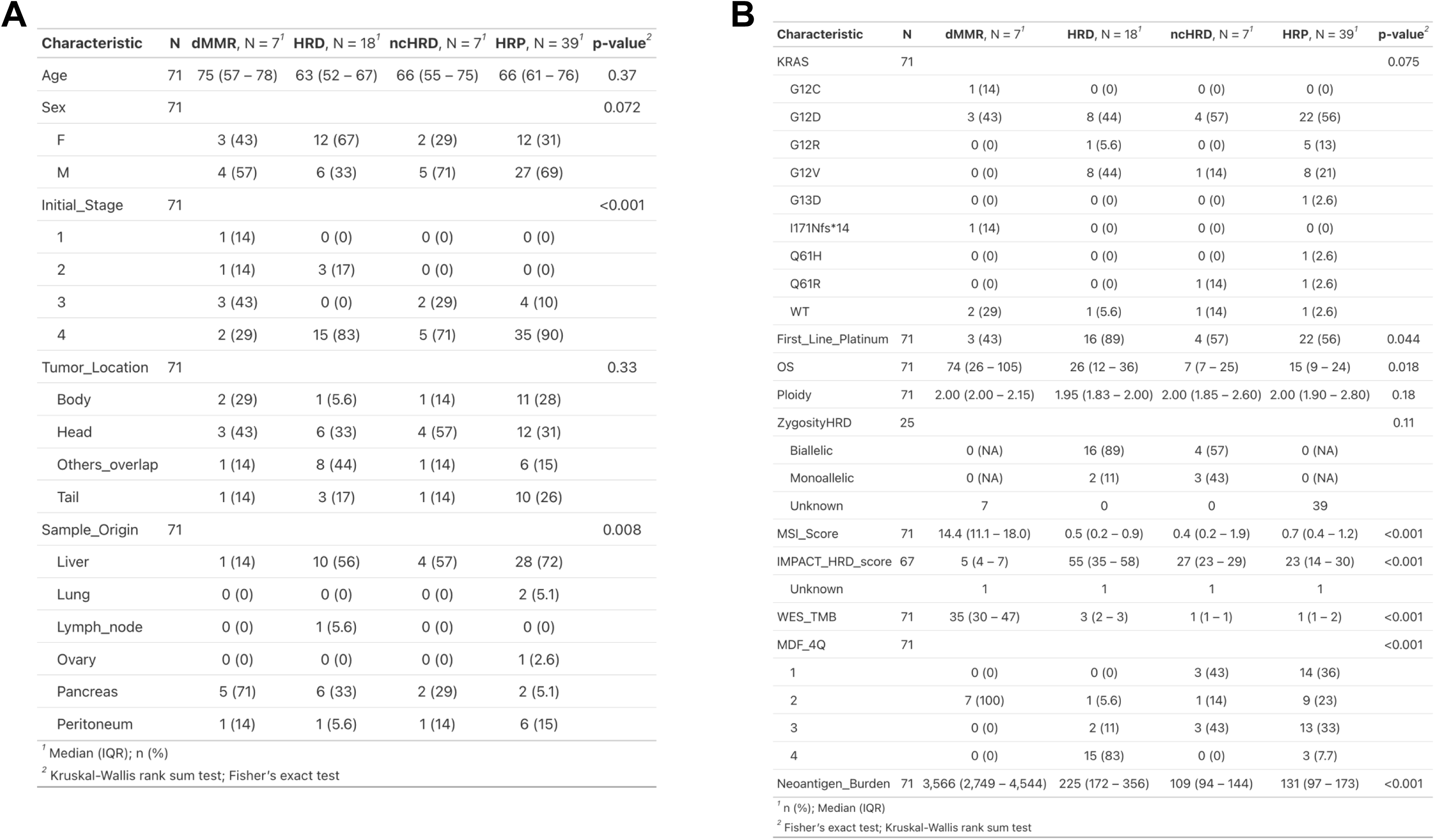
Clinical, molecular and survival characteristics of immunogenic pancreatic cancer stratified by DNA repair deficient subgroups. **(A–B)** Clinical and genomic features across WES-profiled patients (N = 71) stratified by DNA repair status: mismatch repair–deficient (dMMR), homologous recombination–deficient (HRD), non-core HRD (ncHRD), and homologous recombination–proficient (HRP). **(A)** Demographic and clinicopathologic characteristics including age, sex, stage, tumor location, and sample origin. **(B)** Molecular metrics including ploidy, whole-genome duplication (WGD), MSIsensor score, zygosity, HRD score (IMPACT-HRD score), tumor mutational burden (TMB), and MMEJ Deletion Footprint (MDF) quartile. **Abbreviations:** PC, pancreatic cancer; dMMR, mismatch repair deficient; HRD, homologous recombination deficient; ncHRD, non-core HRD; HRP, homologous recombination proficient; TMB, tumor mutation burden; FGA, fraction genome altered; ICB, immune checkpoint blockade; NK, natural killer; MMEJ, microhomology-mediated end joining; MDF, MMEJ Deletion Footprint.

**Table S2.**
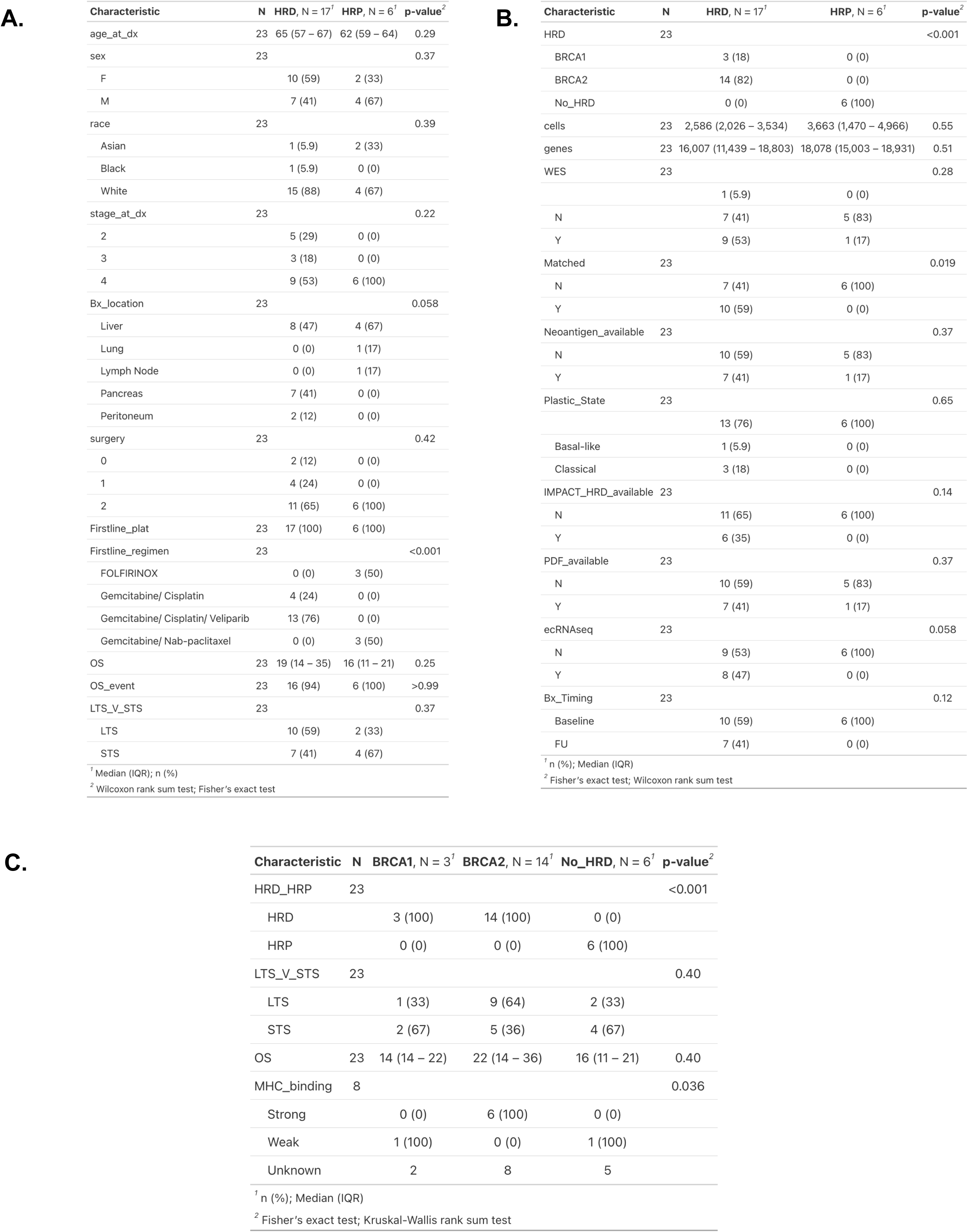
Clinical and Molecular Characteristics of HRD and HRP Pancreatic Tumor Biopsies in the snRNA-seq Cohort. **(A–B)** Baseline demographics, clinical features, and genomic data from the 23 tumors profiled by single-nucleus RNA-seq (HRD=17, HRP=6). **(A)** No significant differences were observed in age, sex, race, tumor stage, biopsy site, or survival outcomes between HRD and HRP tumors. There were more liver biopsy samples HRD tumors were significantly enriched for *BRCA2* mutations and more likely to have received gemcitabine/cisplatin/veliparib as first-line therapy (p<0.001). **(B)** Molecular characteristics showed that HRD tumors more frequently had matched WES data and available neoantigen calls, with a trend toward greater MHC-I binding strength in predicted epitopes (not statistically significant). No differences were observed in cell or gene counts or plasticity state classification. **(C)** Stratified analysis comparing *BRCA1* mutation (N=3), *BRCA2* mutation (N=14), and non-HRD (N=6) tumors.

| Characteristic | N | HRD, N = 17<sup>1</sup> | HRP, N = 6<sup>1</sup> | p-value<sup>2</sup> |
| --- | --- | --- | --- | --- |
| age_at_dx | 23 | 65 (57 – 67) | 62 (59 – 64) | 0.29 |
| sex | 23 |  |  | 0.37 |
| F |  | 10 (59) | 2 (33) |  |
| M |  | 7 (41) | 4 (67) |  |
| race | 23 |  |  | 0.39 |
| Asian |  | 1 (5.9) | 2 (33) |  |
| Black |  | 1 (5.9) | 0 (0) |  |
| White |  | 15 (88) | 4 (67) |  |
| stage_at_dx | 23 |  |  | 0.22 |
| 2 |  | 5 (29) | 0 (0) |  |
| 3 |  | 3 (18) | 0 (0) |  |
| 4 |  | 9 (53) | 6 (100) |  |
| Bx_location | 23 |  |  | 0.058 |
| Liver |  | 8 (47) | 4 (67) |  |
| Lung |  | 0 (0) | 1 (17) |  |
| Lymph Node |  | 0 (0) | 1 (17) |  |
| Pancreas |  | 7 (41) | 0 (0) |  |
| Peritoneum |  | 2 (12) | 0 (0) |  |
| surgery | 23 |  |  | 0.42 |
| 0 |  | 2 (12) | 0 (0) |  |
| 1 |  | 4 (24) | 0 (0) |  |
| 2 |  | 11 (65) | 6 (100) |  |
| Firstline_plat | 23 | 17 (100) | 6 (100) |  |
| Firstline_regimen | 23 |  |  | <0.001 |
| FOLFIRINOX |  | 0 (0) | 3 (50) |  |
| Gemcitabine/ Cisplatin |  | 4 (24) | 0 (0) |  |
| Gemcitabine/ Cisplatin/ Veliparib |  | 13 (76) | 0 (0) |  |
| Gemcitabine/ Nab-paclitaxel |  | 0 (0) | 3 (50) |  |
| OS | 23 | 19 (14 – 35) | 16 (11 – 21) | 0.25 |
| OS_event | 23 | 16 (94) | 6 (100) | >0.99 |
| LTS_V_STS | 23 |  |  | 0.37 |
| LTS |  | 10 (59) | 2 (33) |  |
| STS |  | 7 (41) | 4 (67) |  |
| <sup>1</sup> Median (IQR); n (%) |  |  |  |  |
| <sup>2</sup> Wilcoxon rank sum test; Fisher’s exact test |  |  |  |  |

B.
| Characteristic | N | HRD, N = 17<sup>1</sup> | HRP, N = 6<sup>1</sup> | p-value<sup>2</sup> |
| --- | --- | --- | --- | --- |
| HRD | 23 |  |  | <0.001 |
| BRCA1 |  | 3 (18) | 0 (0) |  |
| BRCA2 |  | 14 (82) | 0 (0) |  |
| No_HRD |  | 0 (0) | 6 (100) |  |
| cells | 23 | 2,586 (2,026 – 3,534) | 3,663 (1,470 – 4,966) | 0.55 |
| genes | 23 | 16,007 (11,439 – 18,803) | 18,078 (15,003 – 18,931) | 0.51 |
| WES | 23 |  |  | 0.28 |
|  |  | 1 (5.9) | 0 (0) |  |
| N |  | 7 (41) | 5 (83) |  |
| Y |  | 9 (53) | 1 (17) |  |
| Matched | 23 |  |  | 0.019 |
| N |  | 7 (41) | 6 (100) |  |
| Y |  | 10 (59) | 0 (0) |  |
| Neoantigen_available | 23 |  |  | 0.37 |
| N |  | 10 (59) | 5 (83) |  |
| Y |  | 7 (41) | 1 (17) |  |
| Plastic_State | 23 |  |  | 0.65 |
|  |  | 13 (76) | 6 (100) |  |
| Basal-like |  | 1 (5.9) | 0 (0) |  |
| Classical |  | 3 (18) | 0 (0) |  |
| IMPACT_HRD_available | 23 |  |  | 0.14 |
| N |  | 11 (65) | 6 (100) |  |
| Y |  | 6 (35) | 0 (0) |  |
| PDF_available | 23 |  |  | 0.37 |
| N |  | 10 (59) | 5 (83) |  |
| Y |  | 7 (41) | 1 (17) |  |
| ecRNAseq | 23 |  |  | 0.058 |
| N |  | 9 (53) | 6 (100) |  |
| Y |  | 8 (47) | 0 (0) |  |
| Bx_Timing | 23 |  |  | 0.12 |
| Baseline |  | 10 (59) | 6 (100) |  |
| FU |  | 7 (41) | 0 (0) |  |
| <sup>1</sup> n (%); Median (IQR) |  |  |  |  |
| <sup>2</sup> Fisher’s exact test; Wilcoxon rank sum test |  |  |  |  |

Table S2. Clinical and Molecular Characteristics of HRD and HRP Pancreatic Tumor Biopsies in the snRNA-seq Cohort
| Characteristic | N | BRCA1, N = 3<sup>1</sup> | BRCA2, N = 14<sup>1</sup> | No_HRD, N = 6<sup>1</sup> | p-value<sup>2</sup> |
| --- | --- | --- | --- | --- | --- |
| HRD_HRP | 23 |  |  |  | <0.001 |
| HRD |  | 3 (100) | 14 (100) | 0 (0) |  |
| HRP |  | 0 (0) | 0 (0) | 6 (100) |  |
| LTS_V_STS | 23 |  |  |  | 0.40 |
| LTS |  | 1 (33) | 9 (64) | 2 (33) |  |
| STS |  | 2 (67) | 5 (36) | 4 (67) |  |
| OS | 23 | 14 (14 – 22) | 22 (14 – 36) | 16 (11 – 21) | 0.40 |
| MHC_binding | 8 |  |  |  | 0.036 |
| Strong |  | 0 (0) | 6 (100) | 0 (0) |  |
| Weak |  | 1 (100) | 0 (0) | 1 (100) |  |
| Unknown |  | 2 | 8 | 5 |  |
| <sup>1</sup> n (%); Median (IQR) |  |  |  |  |  |
| <sup>2</sup> Fisher’s exact test; Kruskal-Wallis rank sum test |  |  |  |  |  |

## SUPPLEMENTARY

**Figure S1.**
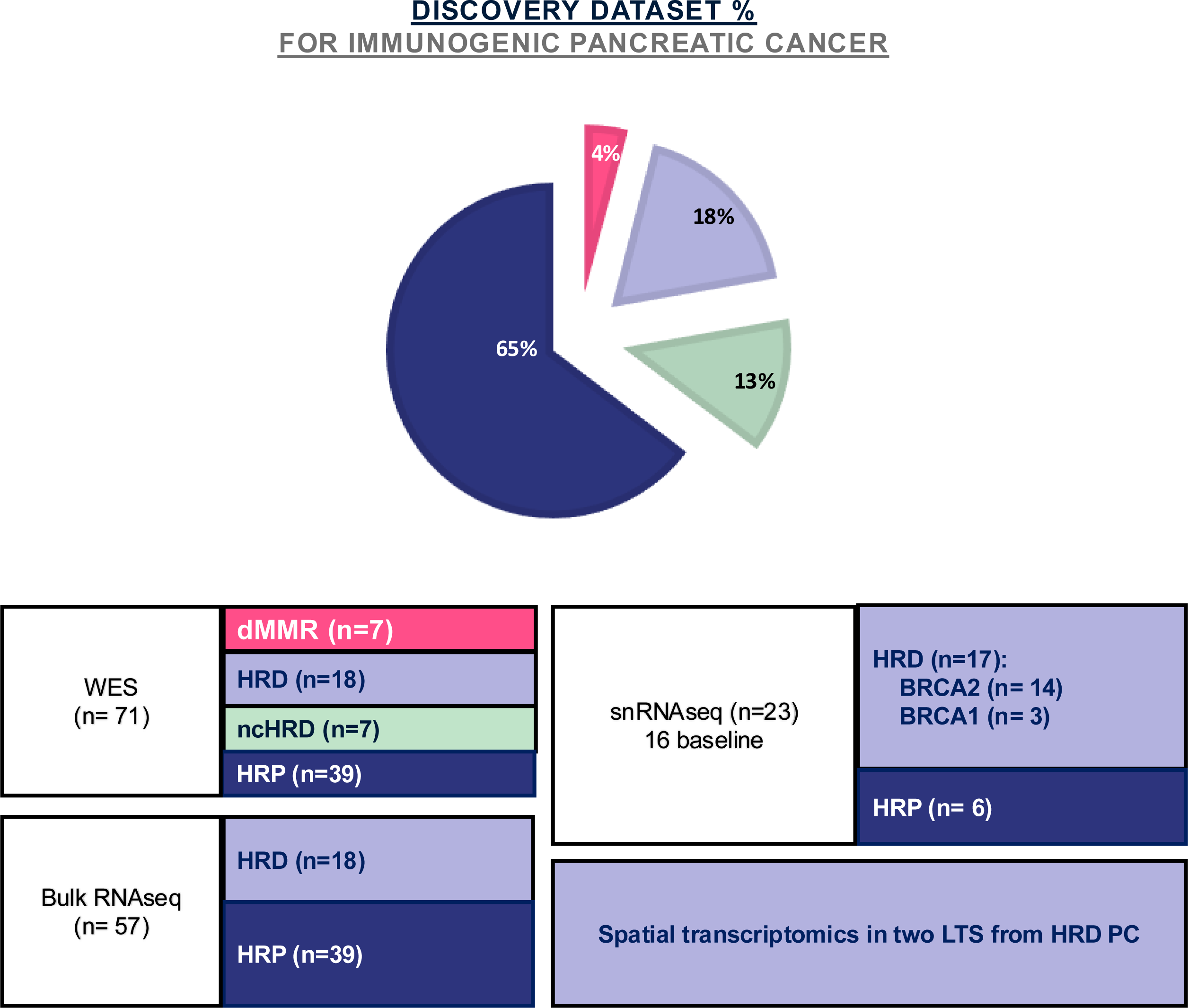
Multiomic profiling overview of the discovery immunogenic pancreatic cancer (iPC) dataset. Schematic summary of the iPC multiomic assay design, depicting data availability (WES, bulk RNA-seq, snRNA-seq, and mIHC/mIF) across DDR-defined subgroups (dMMR, HRD, ncHRD, HRP). Baseline and follow-up (FU) samples, including matched longitudinal sets (n = 5 pairs), are indicated. **Abbreviations:** WES, whole exome sequencing; ecRNAseq, exome-captured RNAseq; snRNAseq, single-nucleus RNAseq; STS, short-term survivor; LTS, long-term survivor; mIHC, multiplxe immunohistochemistry; mIF, multiplex immunofluorescent; PC, pancreatic cancer; dMMR, mismatch repair deficient; HRD, homologous recombination deficient; ncHRD, non-core HRD; HRP, homologous recombination proficient

**Figure S2.**
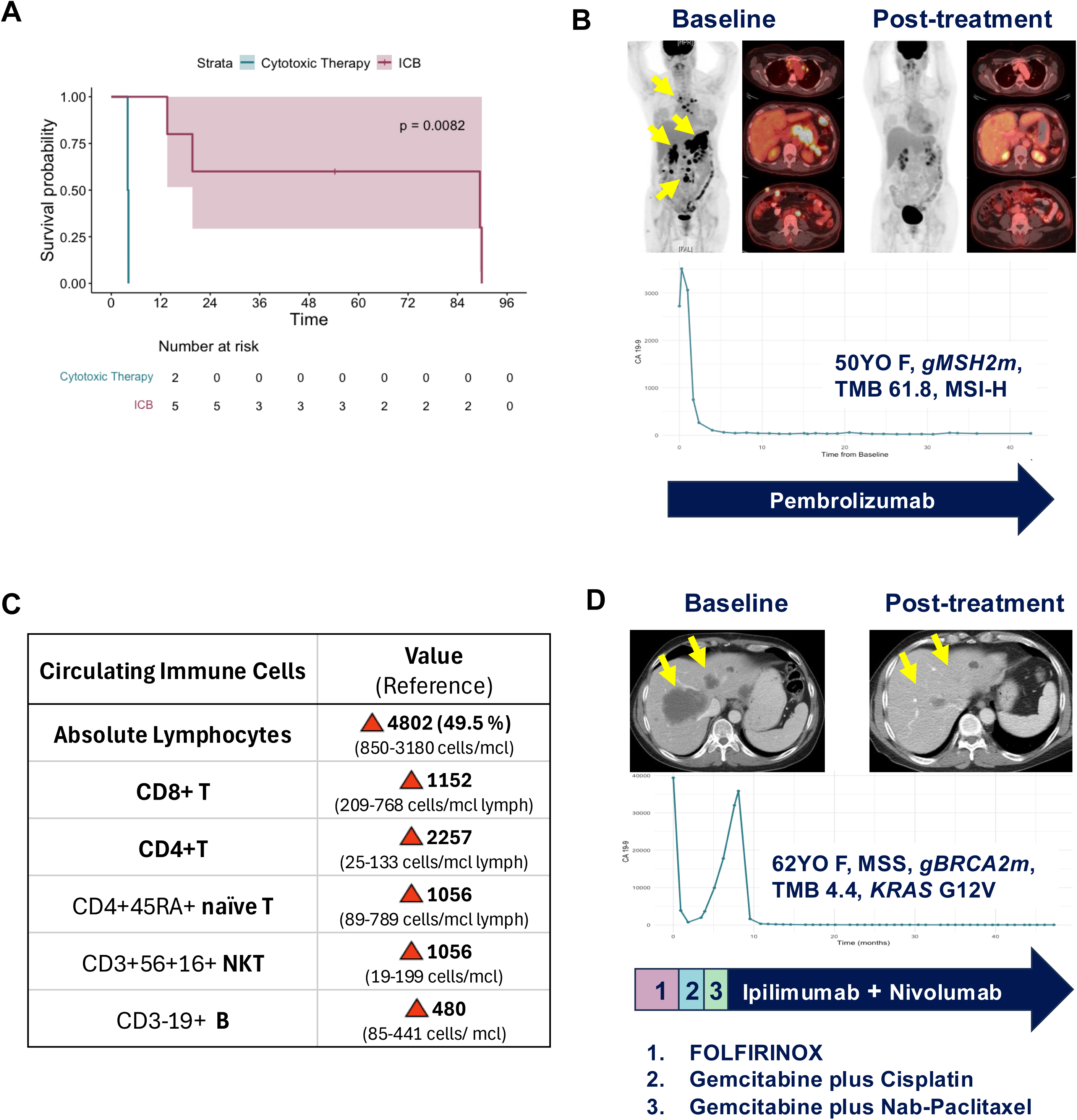
Clinical evidence of immunogenic pancreatic cancer with DNA damage repair deficiency. Mismatch repair deficient (dMMR) pancreatic cancer represents the most clearly immunogenic subset of pancreatic cancer responsive to immune checkpoint blockade (ICB). **(A)** Kaplan–Meier overall survival (OS) curve comparing patients with advanced dMMR pancreatic cancer treated with immune checkpoint blockade (ICB, red) versus cytotoxic chemotherapy (blue) (p=0.0082, N=7). **(B)** PET–CT imaging of an exceptional responder with metastatic pancreatic cancer harboring *gMSH2* mutation, MSI-H status, and high tumor mutational burden (TMB 61.8). Imaging demonstrates complete radiographic response following pembrolizumab therapy. Longitudinal CA19-9 levels show rapid normalization and sustained response during treatment. **(C)** Post-treatment peripheral immune profiling from the same patient demonstrating elevated circulating lymphocytes with increased CD8 ⁺ T cells, CD4⁺ T cells, and CD3⁺CD56⁺CD16⁺ natural killer T (NKT) cells. **(D)** Radiographic response in a patient with metastatic pancreatic cancer harboring germline *BRCA2* mutation (TMB 4.4, MSS) treated with ipilimumab plus nivolumab in the fourth-line setting, illustrating potential immune responsiveness in homologous recombination deficient (HRD) pancreatic cancer.

**Figure S3.**
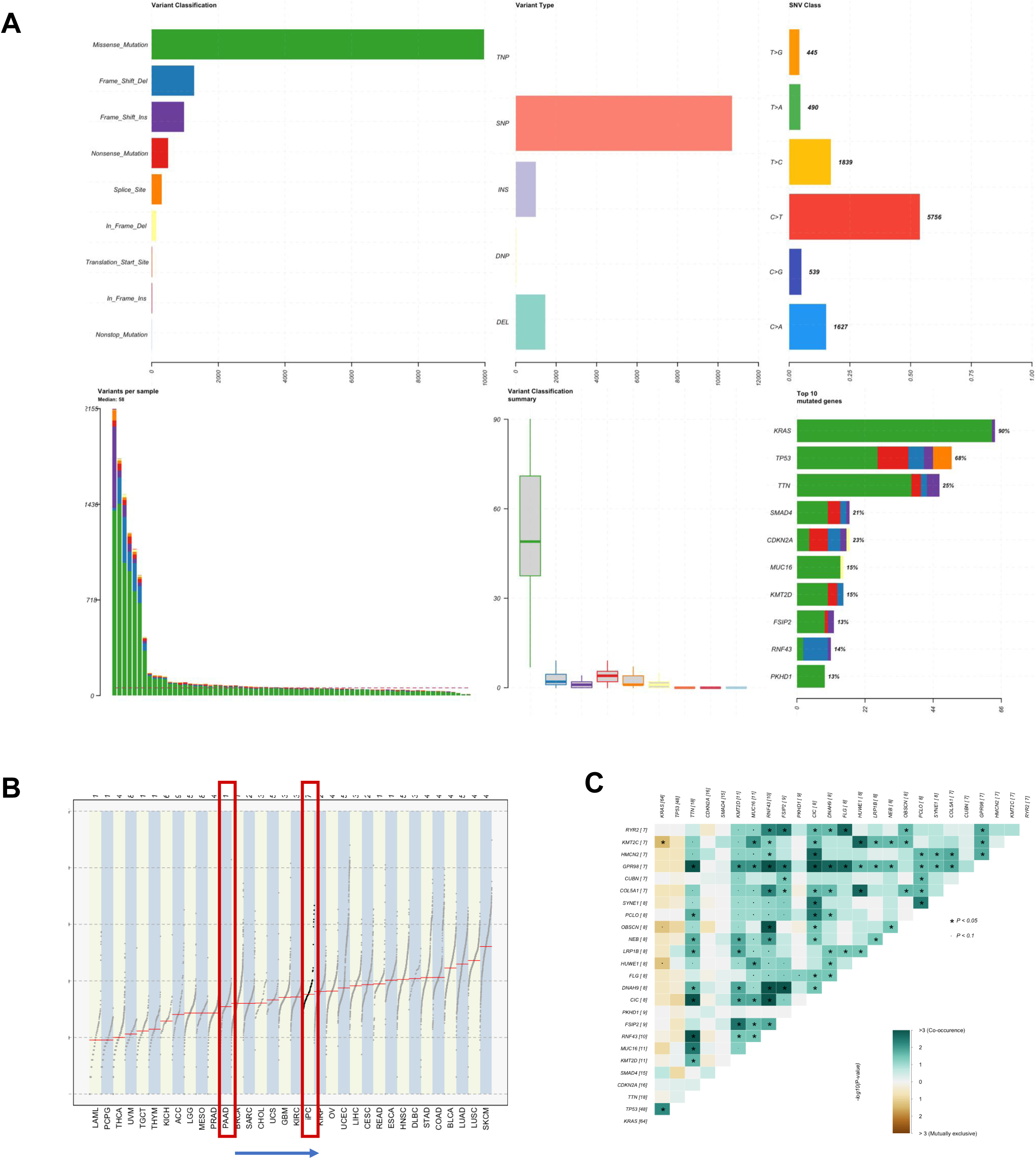
Somatic mutational landscape and frequent non-canonical mutations in DNA repair deficient subgroups. **(A)** Mutational summary of the iPC discovery cohort (N = 73) profiled by whole-exome sequencing. Panels include: **Top left**: Variant classification by type (missense, frameshift, etc.) **Top middle**: Variant type distribution (SNP, INS, DEL) **Top right**: Single-nucleotide variant (SNV) context, dominated by C>T transitions, **Bottom left**: Per-sample mutation count ranked by burden, **Bottom middle**: TMB distribution across samples, **Bottom right**: Most frequently mutated genes across the cohort, with color-coded subgroup contributions **(B)** Tumor mutation burden across tumor types from TCGA, highlighting immunogenic tumors (iPC) with higher median TMB compared to the general representative PDAC (PAAD). **(C)** Co-occurrence matrix of somatic mutations in frequently altered DDR and immune-relevant genes, revealing strong co-mutation patterns beyond canonical DDR genes (e.g., ARID1A, TP53, ZFHX3).

**Figure S4.**
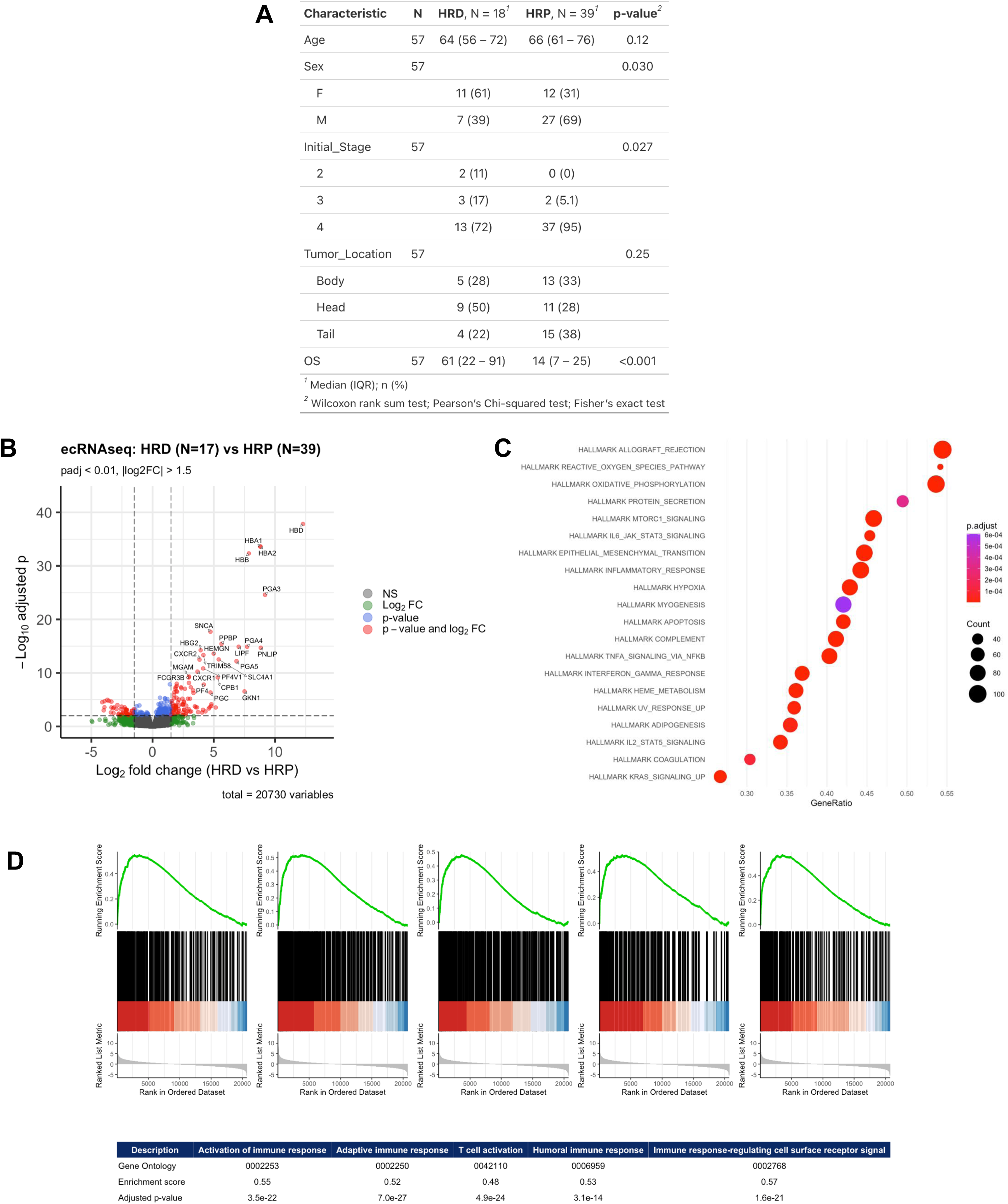
Transcriptomic analysis reveals immunogenicity of homologous recombination deficient pancreatic cancer (HRD PC) (**A)** Demographic and clinical variables including age, sex, stage, tumor location, and overall survival of the ecRNAseq dataset. **(B)** Volcano plot comparing bulk RNA-seq profiles between HRD (N=17) and HRP (N=39) tumors. Differential expression analysis reveals multiple upregulated immune genes in HRD tumors. Threshold: log2 fold-change >1.5 and adjusted p-value<0.01. See STAR Methods: Differential expression analysis. **(C)** Hallmark pathway enrichment analysis (GSEA) identifies immune-related pathways significantly enriched in HRD tumors. See STAR Methods: Hallmark gene set enrichment. **(D)** Gene set enrichment plots for key immune-related GO terms, with associated enrichment scores and adjusted p-values. Top enriched terms include activation of immune response, Adaptive immune response, T cell activation, humoral immune response and immune response-regulating cell surface receptor signal.

**Figure S5.**
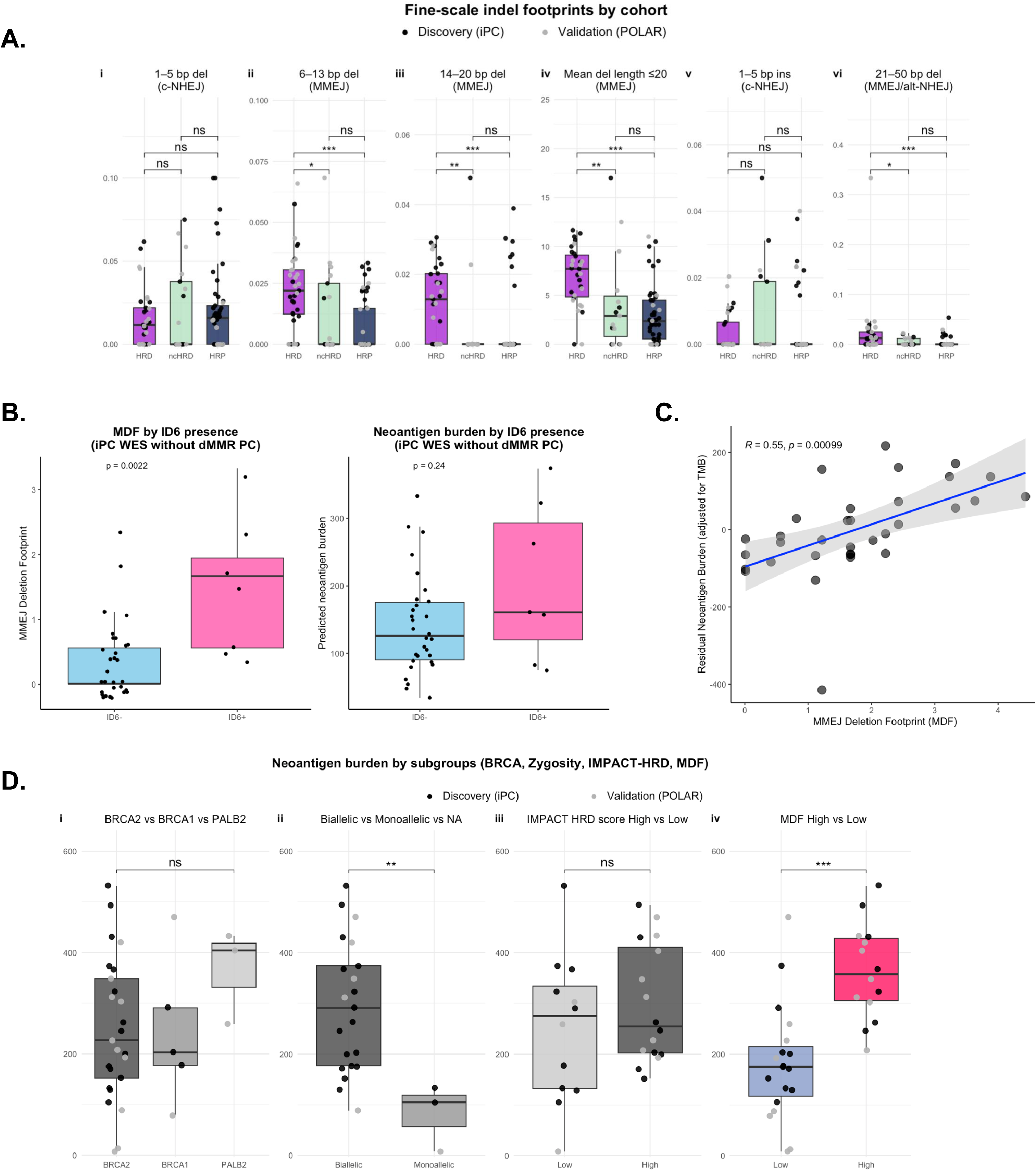
Genomic mutational patterns associated with POLQ-mediated MMEJ activity and neoantigen burden in HRD pancreatic cancer. (A) Fine-scale indel footprint analysis used to derive the MMEJ Deletion Footprint (MDF), which quantifies deletion patterns characteristic of POLQ-mediated microhomology-mediated end joining (MMEJ). HRD umors were enriched for deletions in the 6–20 bp range. Discovery iPC cohort samples are shown as black dots and validation POLAR cohort samples as grey dots. **p-values from Wilcoxon tests; **p < 0.01, *p < 0.001. (B) COSMIC indel signature ID6, a marker of MMEJ repair, was associated with higher MDF scores (p = 0.0022) but not with neoantigen burden (p = 0.24). (C) MDF remained positively associated with neoantigen burden after adjustment for tumor mutational burden (TMB) (Pearson R = 0.55, p = 0.00099). (D) Neoantigen burden across HRD-related genomic stratifications. Neoantigen burden was associated with HRD gene zygosity and MDF score but not with BRCA genotype or IMPACT-HRD score.

**Figure S6.**
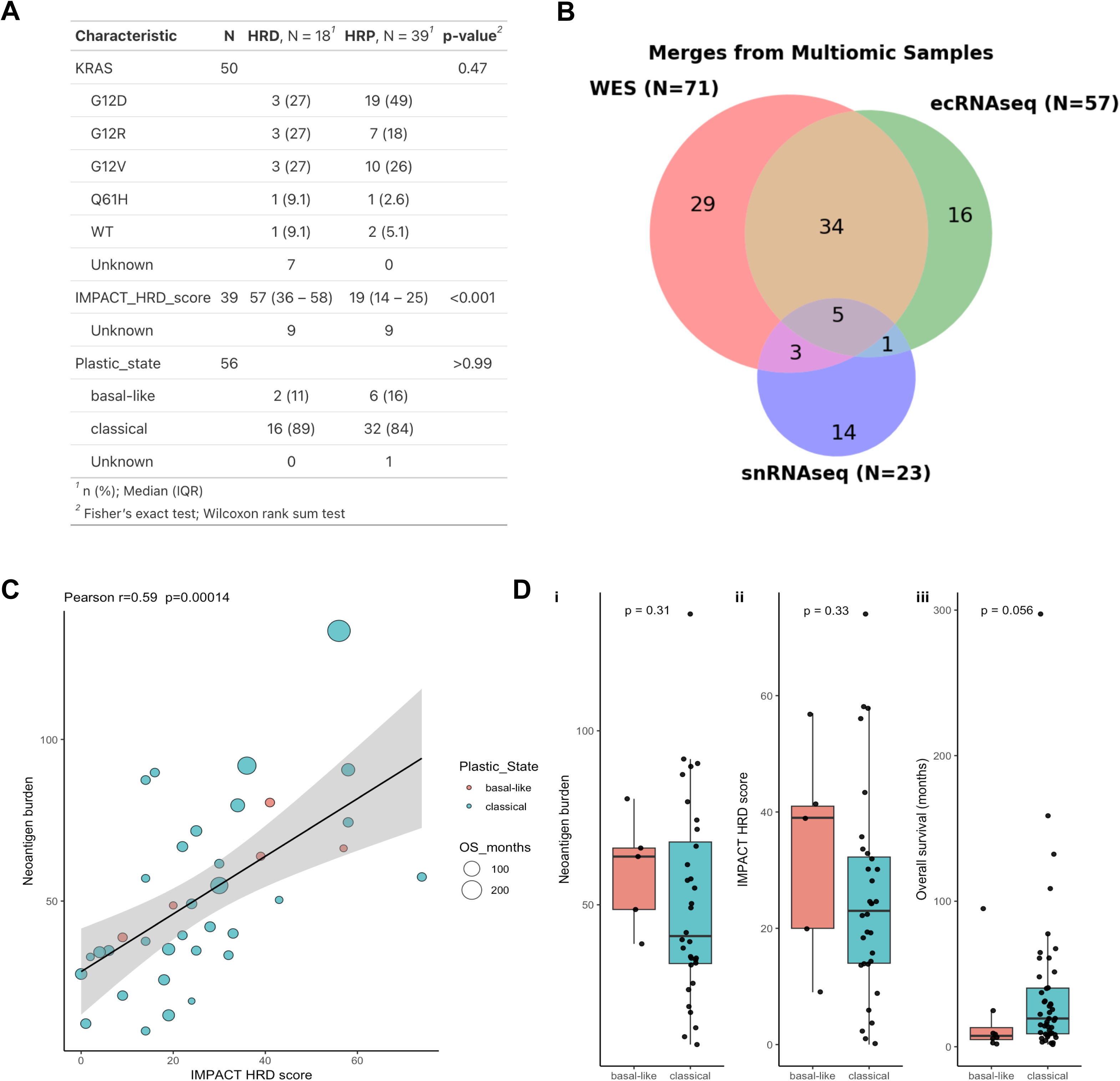
Plastic subtype classification is independent of genomic features of HRD PC. Transcriptional plastic subtype features comparison with clinic-genomic feature: **(A)** KRAS status, IMPACT-HRD score, Plastic state in PurIST available samples (N=56) by HRD and HRP (**B)** Venn Diagram for Merges from multiomic samples **(C)** Neoantigen Burden is associated with HRD score by IMPACT-HRD (R=0.59, p=0.00014). **(D)** Boxplots comparing basal-like and classical subtypes in by: (i) Neoantigen burden (p=0.31), (ii) IMPACT HRD score (p=0.33) and (iii) Overall survival (p=0.056). Classical tumors show longer survival despite comparable HRD score and neoantigen burdens, suggesting that plasticity state adds prognostic value.

**Figure S7.**
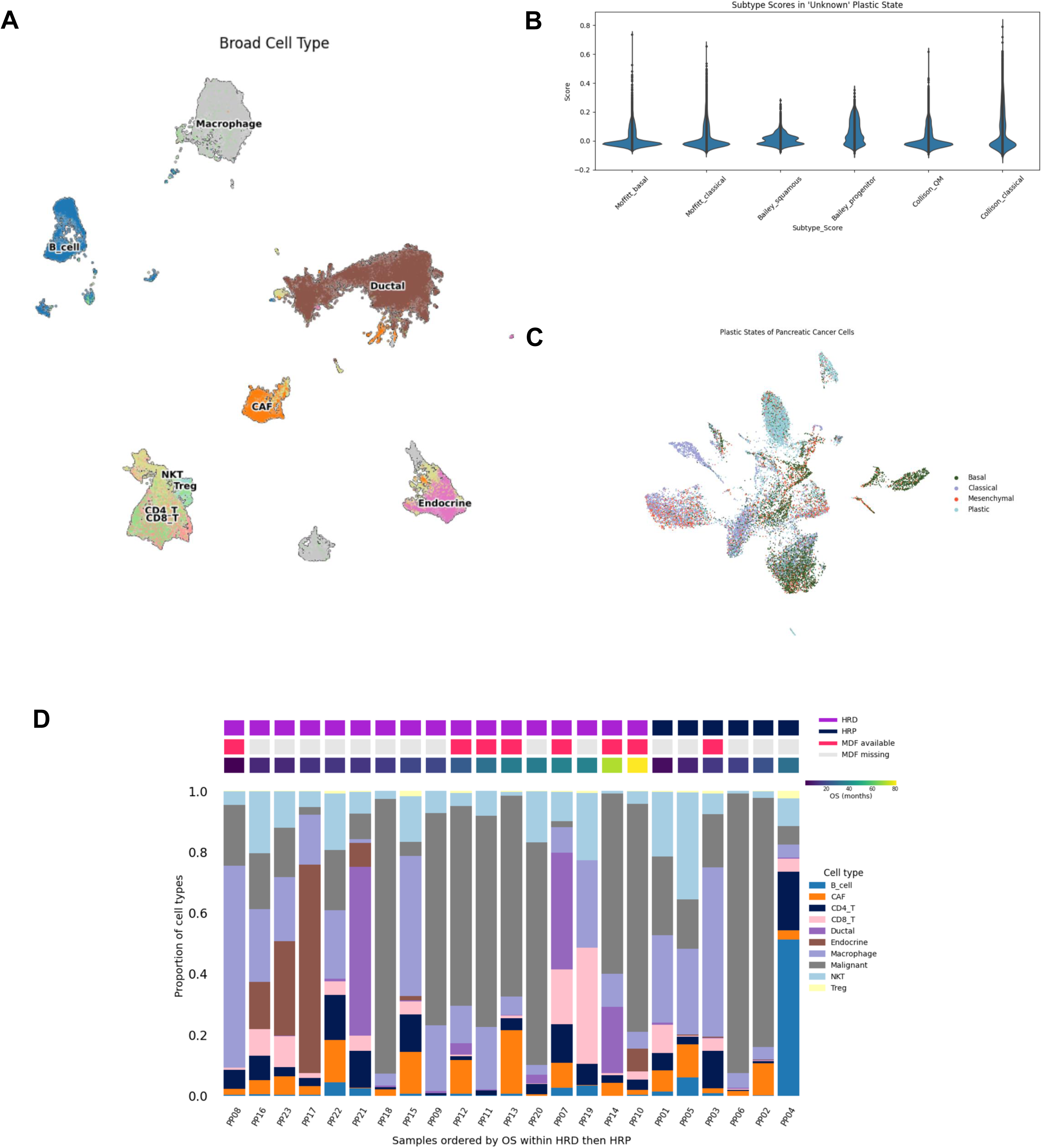
Single nucleus transcriptomics reveals malignant cell plastic states concordant with established pancreatic cancer subtypes. Single-nucleus RNA-seq delineates transcriptional plasticity in pancreatic cancer malignant cells: **(A)** broad cell type annotation across tumor biopsies (N=23), (**B)** malignant cell plastic subtype scoring using gne signatures corresponding to classical, basal-like, progenitor, and quasi-mesenchymal programs, **(C)** UMAP visualization of malignant cell plastic states **(D)** proportional composition of cell types per sample in the increasing order of overall survival and HRD status with annotation bars and MDF availability

**Figure S8.**
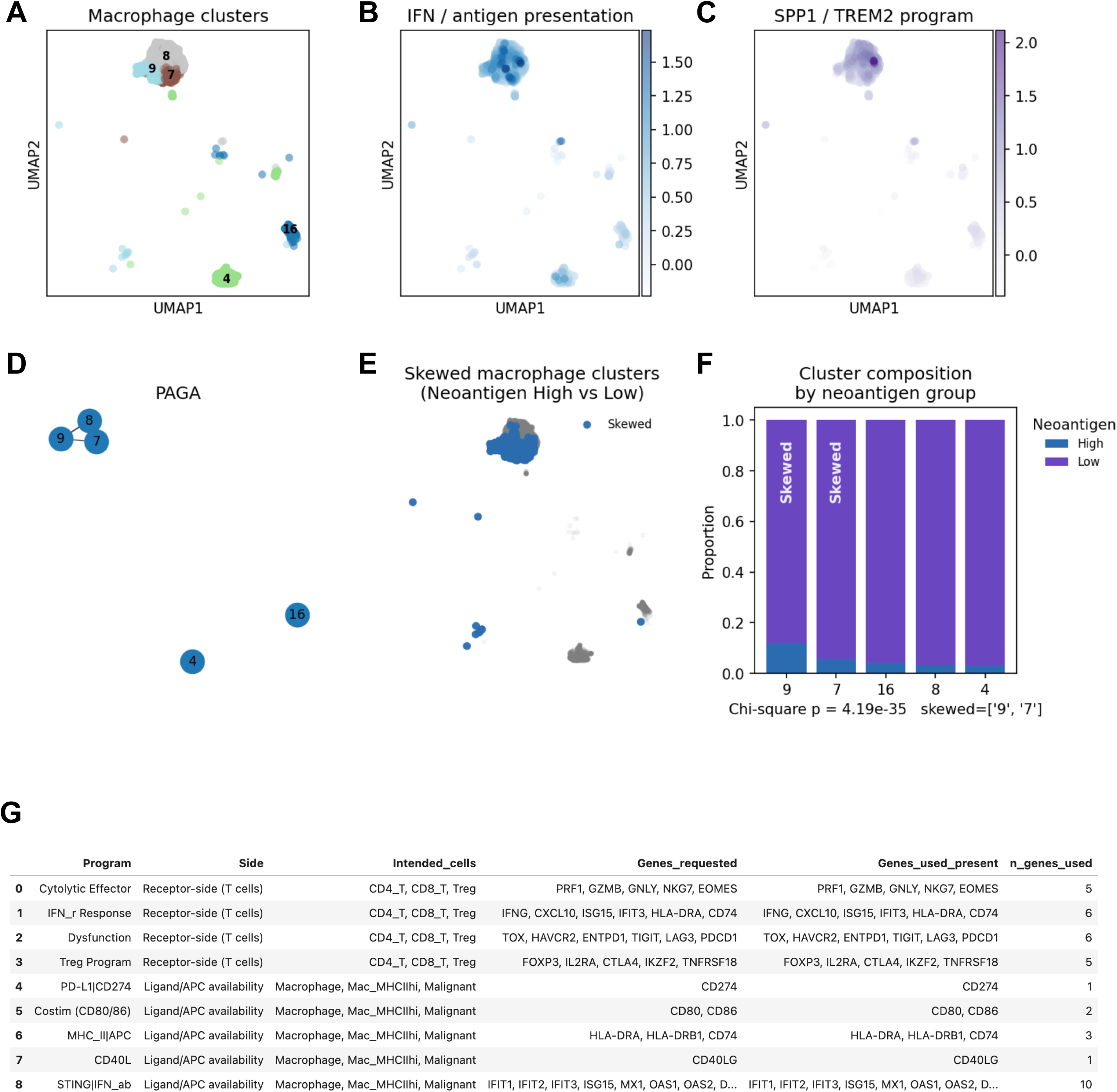
Macrophage clustering, trajectory structure, and immune synapse gene programs. **(A)** UMAP of macrophage clusters identified from the myeloid compartment. **(B)** Module score for interferon and antigen presentation programs across macrophage populations. **(C)** Module score for suppressive TAM programs defined by **SPP1** and **TREM2**. **(D)** PAGA graph showing connectivity among macrophage clusters, illustrating relationships among macrophage states. **(E)** UMAP highlighting macrophage clusters skewed by tumor neoantigen burden (high vs low). **(F)** Cluster composition by neoantigen group (chi-square test shown). **(G)** Gene programs used to quantify immune synapse signaling, including cytolytic T cell, interferon response, dysfunction/Treg programs, and APC-associated ligand programs (PD-L1, CD80/CD86, MHC-II, CD40L, STING/IFN).

